# A two-color haploid genetic screen identifies novel host factors involved in HIV latency

**DOI:** 10.1101/2021.01.20.427543

**Authors:** Michael D Röling, Mahsa Mollapour Sisakht, Enrico Ne, Panagiotis Moulos, Mateusz Stoszko, Elisa De Crignis, Helen Bodmer, Tsung Wai Kan, Maryam Akbarzadeh, Vaggelis Harokopos, Pantelis Hatzis, Robert-Jan Palstra, Tokameh Mahmoudi

## Abstract

To identify novel host factors as putative targets to reverse HIV latency, we performed an insertional mutagenesis genetic screen in a latently HIV-1-infected pseudo-haploid KBM7 cell line (Hap-Lat). Following mutagenesis, insertions were mapped to the genome and bioinformatic analysis resulted in the identification of 69 candidate host genes involved in maintaining HIV-1 latency. A select set of candidate genes was functionally validated using shRNA mediated depletion in latent HIV-1 infected J-Lat A2 and 11.1 T cell lines. We confirmed ADK, CHD9, CMSS1, EVI2B, EXOSC8, FAM19A, GRIK5, IRF2BP2, NF1, and USP15 as novel host factors involved in the maintenance of HIV latency. Chromatin immunoprecipitation assays indicated that CHD9, a Chromodomain Helicase DNA-binding protein, maintains HIV latency via direct association with the HIV 5’LTR, and its depletion results in increased histone acetylation at the HIV-1 promoter, concomitant with HIV-1 latency reversal. FDA-approved inhibitors 5-Iodotubercidin, Trametinib, and Topiramate, targeting ADK, NF1, and GRIK5, respectively were characterized for their latency reversal potential. While 5-Iodotubercidin exhibited significant cytotoxicity in both J-Lat and primary CD4+ T cells, Trametinib reversed latency in J-Lat cells but not in latently HIV-1-infected primary CD4+ T cells. Crucially, Topiramate reversed latency in cell-line models and latently infected primary CD4+ T cells, without inducing T cell activation or significant toxicity. Thus, using an adaptation of a haploid forward genetic screen, we identified novel and druggable host factors contributing to HIV-1 latency.

**Importance:** A reservoir of latent HIV-1-infected cells persists in the presence of combination antiretroviral therapy (cART), representing a major obstacle for viral eradication. Reactivation of the latent HIV-1 provirus is part of curative strategies which aim to promote clearance of the infected cells. Using a two-color haploid screen, we identified 69 candidate genes as latency maintaining host factors and functionally validated a subset of 10 of those in additional T-cell based cell line models of HIV-1 latency. We further demonstrated that CHD9 is associated with HIV-1’s promoter, the 5’LTR while this association is lost upon reactivation. Additionally, we characterized the latency reversal potential of FDA compounds targeting ADK, NF1, and GRIK5 and identify the GRIK5 inhibitor Topiramate as a viable latency reversal agent with clinical potential.

## Introduction

Combination antiretroviral therapy (cART) has proven to effectively abrogate viral replication in HIV-1-infected patients and has substantially reduced AIDS-related mortality. However, cART is not curative and patients must remain on life-long medication regiments, as interruption of the therapy leads to rapid rebound of viral replication (1). This is due to the persistence of a reservoir of latently infected cells, harboring replication-competent provirus blocked at the level of gene expression, that escape clearance by the immune system (2). Therapeutic strategies toward HIV-1 cures aim to inactivate, reduce or completely eradicate the latent reservoir, such that, upon cessation of cART, the patient’s immune system can effectively control the infection or fully clear it (2). Strategies aiming to reduce or eliminate the reservoir rely on drugs, termed latency-reversing agents (LRAs), capable of inducing the latent HIV infected cells to express viral genes to render infected cells susceptible to cytopathic effects and/or recognition and clearance by the immune system (3). Much focus has therefore been placed on finding small molecules for activating transcription of the latent HIV-1 provirus, which are not cytotoxic and do not induce harmful T cell activation or proliferation.

The identification of molecules capable of inducing HIV gene expression has been largely accomplished using candidate approaches, which build on existing knowledge of transcription factors and co-factors that bind to and regulate transcription at the HIV LTR (4–6). One of the most clinically studied classes of molecules in HIV latency reversal, HDAC inhibitors (HDACis) such as SAHA (7), valproic acid (8), rhomidepsin (9) and M344 (10), target HDACs, which have been shown to be recruited to the HIV LTR by multiple transcription factors (11–13) to deacetylate histones and repress transcription. Similarly, agonists of the PKC pathway, such as prostratin and bryostratin, have been studied as activators of the HIV LTR, as they induce nuclear localization and LTR binding of the NFκB (p65/p50) heterodimer, a potent activator of HIV transcription (14, 15). While this candidate approach has led to the identification of druggable targets or potential candidate LRAs, none of the LRAs currently under clinical investigation are capable of strong latency reversal in patients or lead to a reduction in the size of the latent HIV reservoir (16, 17). Thus, to identify more potent and clinically relevant LRAs, it is critical to identify the full repertoire of functionally relevant host factors and pathways that play a role in the maintenance of latency.

Complementary to the approach of targeting candidate LTR-bound transcription factors and complexes for inhibition or activation, other studies have embarked on unbiased screens of small molecule libraries to identify compounds capable of reversing HIV latency (18). Alternatively, recent large-scale unbiased gene knock-out/knock-down screens have been employed to unravel the molecular mechanisms of latent HIV (19–28). RNA interference methods, which rely on the reduction of mRNA expression levels, have been widely used as screening platforms in mammalian cells (29). However, this method has been shown to suffer from serious limitations, including the presence of false-positives due to off-target effects and the persistence of residual gene expression, which results in false negatives (30, 31). For example, several RNAi screens have been performed to identify host cell factors essential for HIV infection and replication (21, 32–34); surprisingly, very little overlap was found between the lists of genes identified in these screens, pointing to the need for other screening methods that achieve complete inactivation of genes. Another large-scale unbiased gene disruption approach uses the precision of CRISPR-Cas9 targeting for complete gene knock-outs (35). A recent screen using a lentiviral sgRNA sub-library targeting nuclear proteins identified MINA53 as a possible latency-promoting gene (LPG) (28). In mammalian cells, functional analysis via forward genetic screens and mutational analysis has largely been hampered due to diploidy, as, in somatic cells, when one copy of a gene essential for a cellular process is inactivated, the second copy often remains active and compensates for that partial loss. Pseudo-haploid screens are based on KBM7s, a chronic myelogenous leukemia (CML) cell line, which is haploid for all chromosomes except for chromosome 8 and a 30Mb stretch of chromosome 15 (36), allowing for forward genetics in mammalian cells (37). Using Gene-Trap (GT) retrovirus-mediated mutagenesis for generating a library of gene knockouts in KBM7 cells enables unbiased loss-of-function screening in mutant cells for the identification of host genes essential to a specific cell function. Haploid screens have proven to be a powerful method to identify genes involved in drug import (38, 39), druggable target genes to treat cancer (40–42), key components of cellular pathways (43–47), and genes involved in the pathogenesis of various viruses (48–55) and susceptibility to toxins (37, 56, 57).

To identify novel host genes that could potentially serve as molecular targets for HIV latency reversal, we performed insertional mutagenesis in latently HIV-1-infected KBM7 cells. First, using a flow cytometry activated cell sorting-based strategy described previously (58, 59), we generated a latent pseudo-haploid KBM7/(Hap-Lat) cell line that harbors an integrated transcriptionally silent HIV-1 5’LTR controlling the expression of a GFP reporter gene. Hap-Lat cells were then subjected to GT insertional mutagenesis, using a mCherry reporter GT virus, and Hap-Lat cells expressing both GFP (as a reporter of HIV-1 expression) and mCherry (confirming the presence of a GT integration within an active gene) were then sorted using fluorescence-activated cell sorting (FACS). GT integration sites within these two populations were then amplified by inverse PCR and mapped to the genome by high-throughput sequencing. Computational candidate identification resulted in the identification of 69 candidate genes, whose function is required for maintenance of HIV latency, but which are not essential to cell survival. For functional follow-up experiments, we selected a subset of 16 GT-identified putative HIV latency re-enforcing target genes, which are also expressed in primary CD4+ T cells, and examined the effect of their depletion via shRNA knock-down in maintaining latency in J-Lat T cell lines A2 and 11.1. Functional validation identified 10 putative novel latency promoting genes: ADK, CHD9, CMSS1, GRIK5, USP15, IRF2BP2, EXOSC8, NF1, EVI2B, and FAM19A. Because of the obvious mechanistic potential of CHD9, a Chromodomain Helicase DNA-binding protein, we explored its association with the 5’LTR using ChIP-qPCR analysis. We found CHD9 to be enriched at the 5’LTR in the latent state but to dissociate from the HIV-1 promoter after PMA treatment. Loss of CHD9 enrichment upon re-activation was accompanied by a relative increase in H3 acetylation at the HIV-1 LTR, indicating a functional shift in chromatin organization from repressed to transcriptionally active. We also found three genes in our candidate list, ADK, GRIK5, and NF1 to be targetable by existing small molecule inhibitors. The latency reversal potential of these compounds was evaluated in J-Lat T cell lines A2 and 11.1, and in a primary cell model of HIV latency. We found 5-Iodotubercidin, which inhibits ADK, not to be a viable LRA due to its toxicity in both A2, 11.1, and primary CD4+ T-cells. While the NF1 inhibitor Trametinib reactivated latent HIV in the J-Lat cell lines, it was not effective in the primary cell model of latency. On the other hand, the GRIK5 inhibitor Topiramate reversed HIV-1 latency in both J-Lat T-cell lines as well as in primary CD4+ T cells harboring latent HIV-1, without significant associated cytotoxicity or T cell activation, and thus presents an interesting potential novel LRA.

## Results

### Establishment of a latent pseudo-haploid cell line

To identify potentially druggable host genes as putative molecular targets for HIV latency reversal, we generated pseudo-haploid latent HIV-1-infected cells in which we performed insertional mutagenesis according to a strategy schematically depicted in Figure 1A. A latent HIV-1-infected pseudo-haploid KBM7 cell line was generated according to a previously described strategy used in Jurkat cells (58, 59) (Figure 1B). Subsequently, haploid latent (Hap-Lat) cells harboring a latent integrated HIV-1-derived virus containing a GFP reporter, were subjected to insertional mutagenesis using gene trap virus carrying an mCherry reporter. Instead of using a lethality-selecting scheme, our system relies on fluorescence activated cell sorting (FACS) to select for reactivated cells, marked by elevated GFP expression resulting from insertional mutagenesis of genes essential for maintenance of HIV latency. Cells expressing both GFP and mCherry were FACS-sorted and expanded for multiple rounds, after which GT insertion sites were mapped and identified by inverse PCR and high-throughput sequencing. To establish a latent HIV infection in the pseudo haploid KBM7 system, near-haploid KBM7 cells were infected at low MOI with the single infectious cycle HIV-derived virus LTR-GFP-LTR, in which GFP reporter expression is controlled by the activity of the HIV-1 promoter or 5’LTR (Figure 1A). After infection, the population of GFP negative cells comprising mainly uninfected cells and putative latently infected cells were sorted by FACS and subsequently stimulated with 5ng/μl TNF-α and 350μM Vorinostat. In response, a small percentage of existing latent HIV-infected cells were transcriptionally reactivated and expressed GFP. These GFP positive cells were -sorted by FACS as single cells and expanded (Figure 1B). The resulting clonal latent KBM7 haploid lines were characterized by flow cytometry to determine GFP expression at basal and stimulated states. From the clonal lines established, Hap-Lat#1 was selected for low basal activity and significant reactivation upon stimulation (Figure 1C). To ensure maintenance of haploidy, Haplat#1 cells were periodically sorted to enrich for the 5% of smallest cells (Supplemental figure 1A).

**Figure 1.**
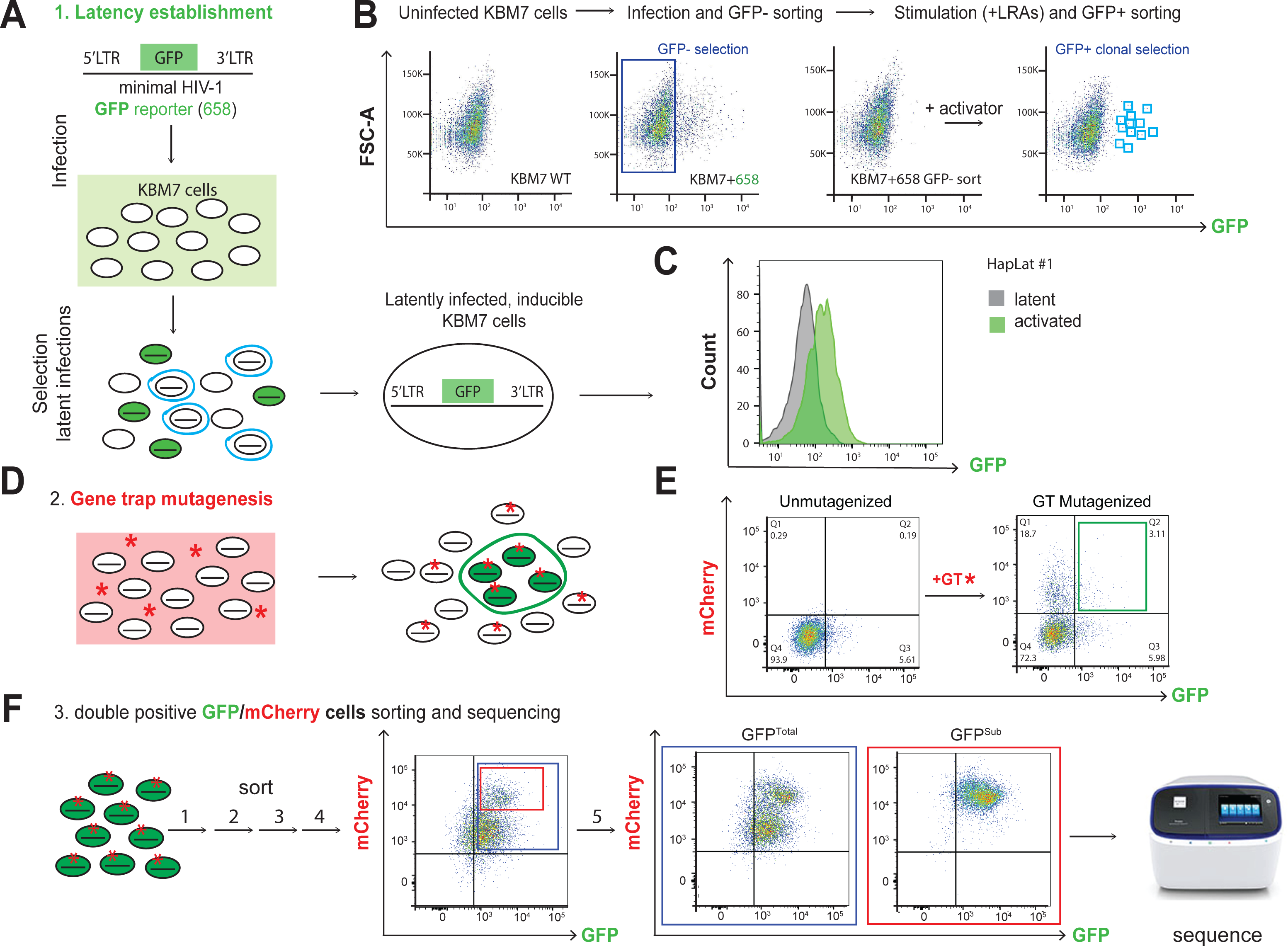
Schematic representation of the two-color haploid screening strategy for identification of novel host factors and cellular pathways involved in the maintenance of HIV latency. (A) Scheme depicting the generation of a clonal latent haploid KBM7 cell line. To this end, haploid KBM7 cells were infected with a minimal HIV virus harboring GFP (HIV-1 658). After stimulation, reactivated cells were sorted and left to expand and revert to a latent state. (B) FACS plots depicting the establishment of haploid HIV-1 latently infected KBM7 cell-line (Hap Lat #1). Parental KBM7 cells were infected with a minimal HIV virus carrying a GFP reporter (HIV-1 658). GFP negative cells, consisting of uninfected and latently infected cells, were sorted by FACS. The polyclonal cell pool was stimulated with a cocktail of latency reversal agents (LRAs) and reactivated cells were clonally sorted by FACS and expanded to generate haploid latent cell lines. (C) Hap-Lat #1 displays low basal activity (GFP expression) but is effectively reactivated using LRAs. (D) Hap-Lat#1 cells were mutagenized by infection with a Gene Trap (GT) virus infection harboring a mCherry reporter. Cells infected with the GT reporter will be mCherry positive (red asterisk). Latently infected KBM7 cells that reactivate following GT mutagenesis will be double positive for GFP and mCherry (Green cells, red asterisk). (E) Representative FACS plots demonstrating gating strategy for sorting double positive cells (GFP, mCherry). (F) Double positive cells (GFP, mCherry) are sorted in multiple rounds to eliminate cells stochastically reverting back to a GFP negative state. During these rounds of sorting, a stable and distinct double positive sub-population (GFP Sub) appears which was sorted separately.

### Gene-trap mutagenesis of Hap-Lat cells

Hap-Lat#1 cells were mutagenized by infection with a murine stem cell virus (MSCV)-derived viral gene-trap (GT) vector containing an inactivated 3’ LTR, an adenoviral splice-acceptor site, an mCherry reporter cassette and a polyA terminator tail (37). Dendritic and myeloid cells are notoriously refractory to retroviral infection (60, 61). We indeed found that infectivity of KBM7 is poor compared to T-cell-derived cell lines SupT1 and Jurkat, as well as other myeloid-derived cell lines (Supplemental Figure 1B). SAMHD1, a nucleotide scavenger, has been identified as a causative restricting host factor that limits the free available pool of nucleotides for reverse transcription (62). To bypass SAMHD1-mediated restriction, we supplemented cells with 2μM nucleosides (dNs), which increased GT infectivity by approximately two-fold (Supplemental figure 1C). GT preferentially inserts in the 5’ regions of genes (63), effectively knocking out gene expression by truncating the native transcript (Supplemental figure 1D). For a full-scale mutagenesis experiment, approximately 200 million Hap-Lat #1 cells were mutagenized using two rounds of infection with GT-mCherry in the presence of exogenously supplied nucleosides. Infection of Hap-Lat#1 with GT-mCherry effectively caused reactivation of a subpopulation of latent KBM7 cells (Figure 1D-E). We reasoned that insertional mutagenesis in genes essential for maintenance of HIV latency would result in Hap-Lat#1 cells expressing the GFP reporter. We determined GT integrations in individual GFP/mCherry double-positive clones and estimated that the sequential infection resulted in 1 to 4 GT integrations per cell, with the majority of cells containing one integration (data not shown). By gating conservatively, approximately 1-4% GFP/mCherry double-positive cells were then sorted and expanded (Figure 1E). After expansion, reactivated cells tend to revert to a latent state (Supplemental figure 1E). To enrich for a more stable GFP-expressing, mCherry GT-containing double-positive cell population, cell sorting was repeated for multiple rounds (Figure 1F). Sequential rounds of sorting led to the appearance of a stable subpopulation within the total double-positive population expressing high levels of mCherry (Figure 1F). To examine any potential biological differences between the two, we separately sorted the total double-positive population and the mCherry-high subpopulation, which we designated GFP^Total^ and GFP^Sub^ respectively, for a final (5^th^) round (Figure 1F). Genomic DNA extracted from GFP^Total^ and GFP^Sub^ obtained in the 5^th^ round of sorting was used to determine GT integrations, while that of a pool of unsorted GT-infected cells was used as a reference.

### Mapping of insertion sites to identify host factors maintaining HIV latency

To determine the host sequences flanking the GT insertion sites, inverse PCR with primers annealing to internal sequences in the gene trap vector followed by amplification was performed. The amplified products were processed for high-throughput sequencing (Figure 1F). For GFP^Sub^ 2 biological replicates, samples A and B, were generated. For GFP^Total^ 3 biological replicates were generated, samples C, D and E. To estimate the sampling depth of our GT, we re-sequenced GFP^Sub^ sample B and GFP^Total^ sample D at greater depth. The resulting NGS datasets were processed for candidate gene identification. A previously described method to analyze GT data, HaSAPPy (Haploid Screen Analysis Package in Python), was rigorously re-implemented, appended with additional steps for quality control, library normalization, and optimized resolution for the selection of integration sites (64). HaSAPPY assigns a Local Outlier Factor (LOF) score to each gene in a sample based on a triplet score derived from the number of putative GT integrations in the sample compared to the reference. For each population, we compiled all genes with an LOF score >3 from each replicate and obtained 686 hits for GFP^Total^ and 382 hits for GFP^Sub^. 183 genes were common to both populations (Supplementary figure S2A). Next, we investigated any potential biological basis for the difference between GFP^Total^ and GFP^Sub^. Since expression levels of the integrated mCherry reporter appear to be on average higher in the GFP^Sub^ population than in the GFP^Total^ population, we wondered if expression levels of the targeted genes were higher in GFP^Sub^. We obtained recently published KBM7 gene expression data (65) and found no substantial difference in the average level of expression between the two populations (Supplementary figure S2B). To determine if there was any difference in the functionality of the GT target genes found in GFP^Total^ and GFP^Sub^, we performed enrichment analysis using GO terms and found no substantial differences in enrichment for biological process and molecular function ontologies (Supplementary figure S2C and D). Similarly, both populations contain a comparable fraction (32.7%) of integrations within non-coding or anti-sense genes (Supplementary figure S2E). Finally, we cross-referenced the GT target genes found in GFP^Total^ and GFP^Sub^ populations to the HIV interaction database (https://www.ncbi.nlm.nih.gov/genome/viruses/retroviruses/hiv-1/interactions/) and found that both the GFP^Total^ and the GFP^Sub^ populations contain similar fractions (21.9% and 20.2%, respectively) of genes previously reported to be involved in HIV biology, fractions which are substantially higher than the 7.4% found for the complete list of ENSEMBL genes (Supplementary figure S2F). In order to limit our extensive list of candidate genes for follow up functional validation, we applied more stringent thresholds for each population and defined candidate genes as having an LOF score equal to or greater than 3 in at least 2 biological replicate samples. We thus identified 19 candidate genes in the GFP^Sub^ population and 55 in the GFP^Total^ population (Figures 2A and B and Table 1).

**Figure 2.**
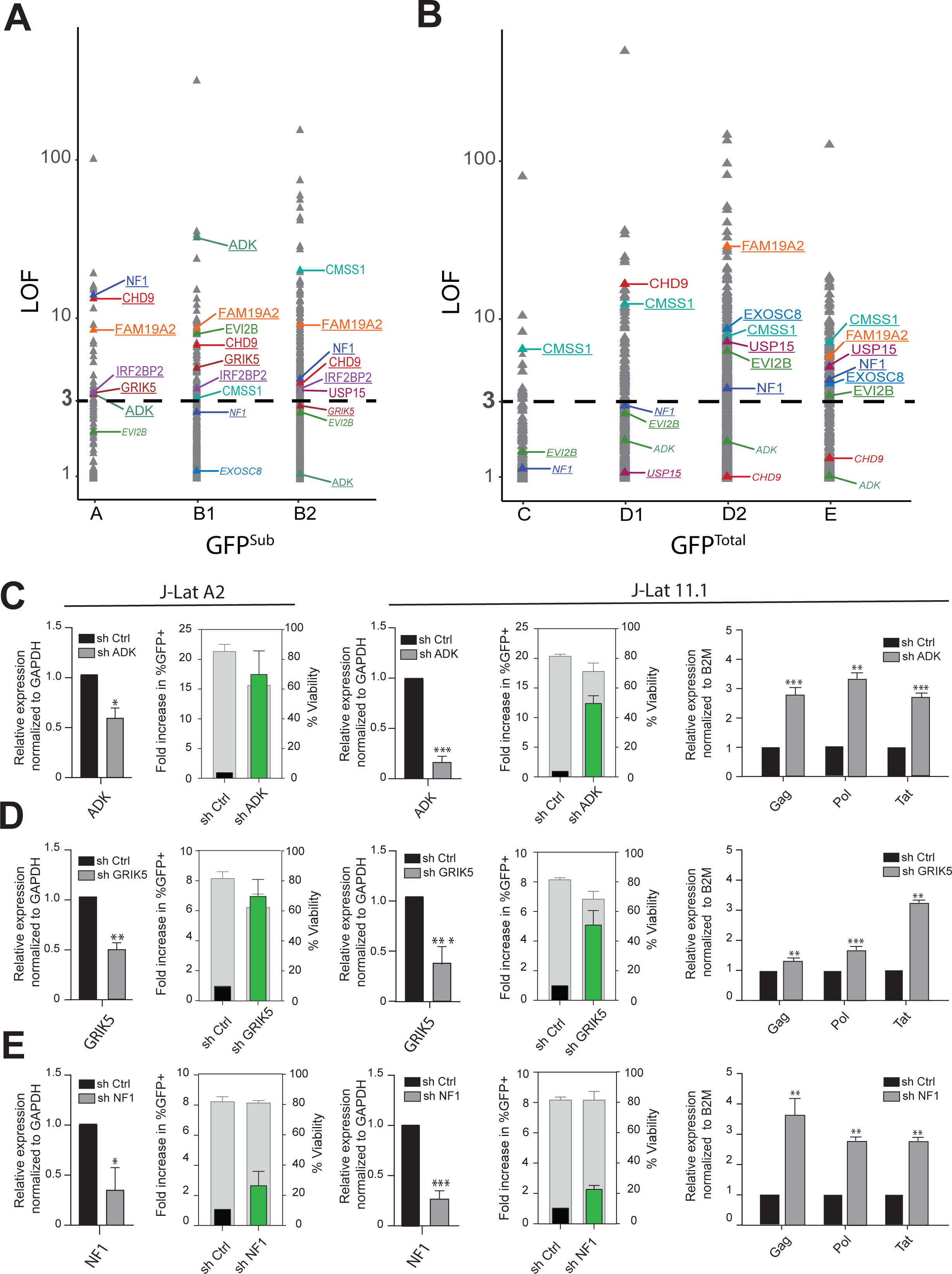
Identification and validation of candidate host factors. (A) LOF scores of genes in the GFP^sub^ population (samples A, B1 and B2), validated candidates are indicated. Genes with LOF>3 are in large font, underlined genes comply to our candidate gene selection criteria and have a LOF>3 in at least two biological replicates within either the GFP^sub^ or GFP^total^ population while genes with LOF<3 but complying to our selection criteria based on other samples are depicted in small font and italics. (B) LOF scores of genes in the GFP^total^ population (samples C,D1,D2 and E), Markings as in Figure 2A. (C) Functional validation of candidate hits ADK, GRIK5, NF1 by shRNA mediated depletion followed by determination of latency reversal by flow-cytometry and RT-PCR in latently infected J-Lat A2 (left panels) and 11.1 cells (right panels). Flow-cytometry bar plots: green bars show the percent of GFP positive cells after knockdown over control (black bar), left y-axis, whereas gray bars show cell viability, right y-axis. Viral reactivation is confirmed by RT-qPCR for viral genes Tat, Gag and Pol in J-Lat 11.1 cells. Statistical significance was calculated using ratio-paired t-test and multiple comparison t-test on Log2 transformed fold changes * – p < 0,05, ** – p < 0,01, *** – p < 0,001,

**Table 1.**
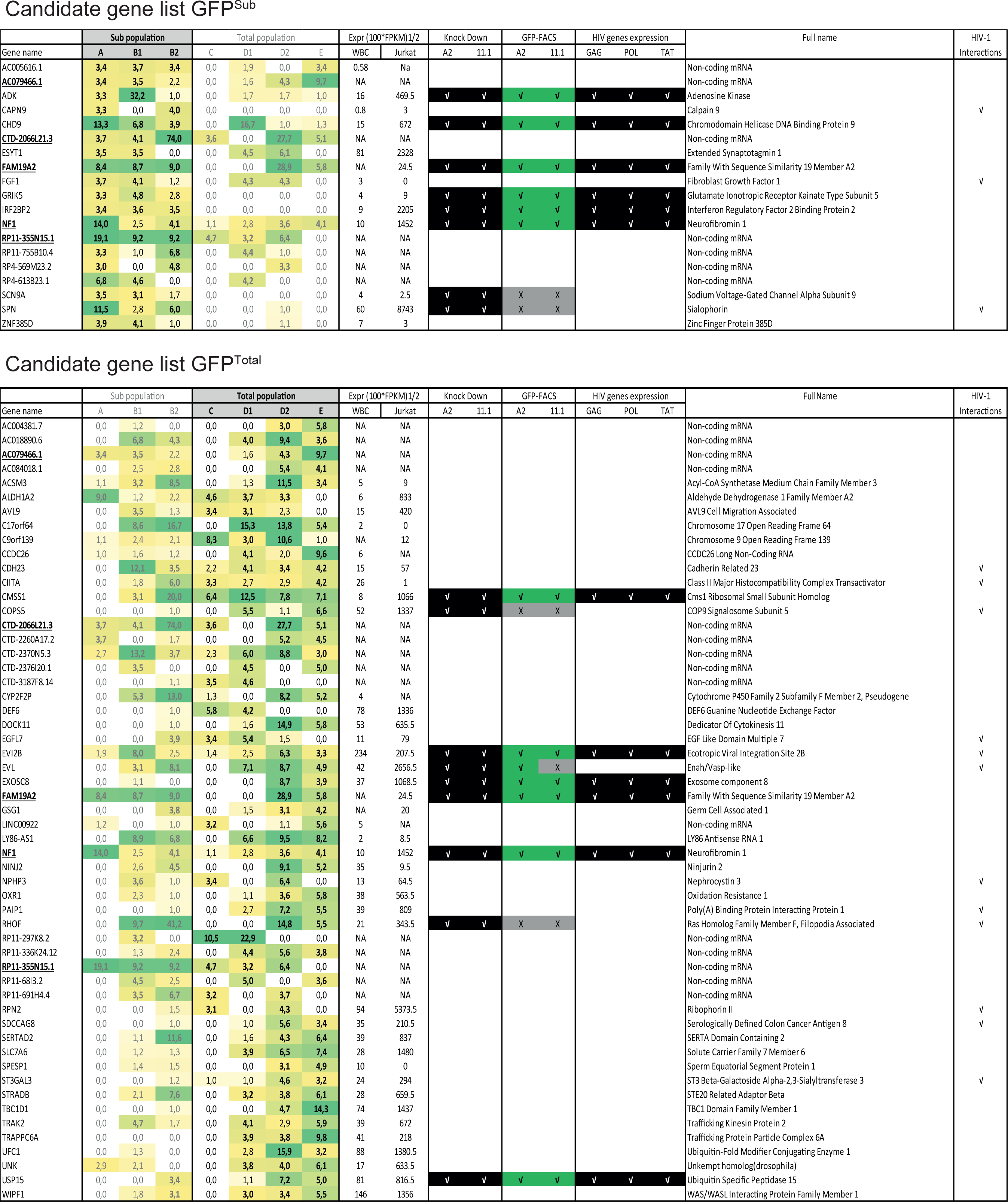
Prioritized candidate list for the GFP^Total^ and the GFP^Sub^ populations. Prioritized candidate genes are defined as having an LOF equal to or greater than 3 in at least 2 independent samples. Overlapping genes are underlined in both lists. Results of shRNA mediated knockdown validation experiments of selected candidate genes in A2 and 11.1 J-Lat cell models are indicated.

### Candidate list validation

Since our bioinformatics analysis did not reveal a defining difference between the GFP^Total^ and the GFP^Sub^ populations, we decided to proceed with functional assays using shRNA-mediated depletion of candidate genes obtained from both populations. To prioritize the candidate genes found in the KBM7 haploid screen for functional validation in the more biologically relevant Jurkat T-cell-based HIV-1 latency models J-Lat A2 and 11.1, we focused on protein-coding genes and took into account LOF scores as well as gene expression in white blood cells (as extracted from the GTEx portal (https://www.gtexportal.org/home/) or Illumina’s Human BodyMap 2.0 project (http://www.ensembl.info/2011/05/24/human-bodymap-2-0-data-from-illumina/)) and the Jurkat T-cell model (66) (Table 1). Eight genes from GFP^Sub^ and nine genes from GFP^Total^ were selected as candidates for validation using shRNA-mediated depletion in J-Lat A2 and J-Lat 11.1 cells. These cells contain a latent HIV-derived GFP reporter driven by the 5’HIV-LTR and are well-established model systems for HIV latency (58, 67). While J-Lat A2 cells contain an integrated latent LTR-Tat-IRES-GFP virus, J-Lat 11.1 cells contain an envelope defective full-length HIV-1 genome expressing GFP in place of Nef. We used flow cytometry to assess reactivation as measured by GFP expression and extracted RNA to assess knockdown of the targeted candidate genes and expression of the HIV genes GAG, POL, and TAT by RT-qPCR (figure 2C-E and Supplemental figure S3 and S4). We observed significant latency reversal upon candidate gene knockdown in 10 of 15 genes depleted by shRNA (ADK, CHD9, CMSS1, EVI2B, EXOSC8, FAM19A, GRIK5, IRF2BP2, NF1, and USP15) in both J-Lat A2 and J-Lat 11.1 cells (Figure 2C-E and Supplemental figure S3 and Table 1). Knockdown of SCN9A, RHOF, SPN, COPS5 and EVL did not result in significant latency reversal in one or both J-Lat models (Supplemental figure S4). These results demonstrate that a significant proportion of the candidate genes found in our myeloid-derived KBM7 haploid screen play a role in maintenance of HIV-1 latency in the more relevant T cell-derived J-Lat A2 and J-Lat 11.1 HIV latency models.

### CHD9 is an LTR-associated repressor of HIV-1 transcription

Interestingly, two genes, CIITA and CHD9, from our candidate list are associated with the GO-term DNA binding (GO:0003677). CIITA is a well-established factor involved in HIV expression and has been previously shown to inhibit Tat function and hence viral replication (68, 69). The Chromodomain helicase DNA binding protein 9 (CHD9) is a member of an ATP-dependent chromatin remodeler family, the members of which modulate DNA-histone interactions and positioning of nucleosomes and play key roles in stem cell regulation, development, and disease (70). Previously, we have shown that chromatin remodeling by another ATP-dependent remodeler, the BAF complex, plays a crucial role in maintenance of HIV-1 latency and its re-activation (59). We therefore decided to further characterize the role of CHD9 in regulating HIV-1 gene expression. We knocked down CHD9 using a lentivirally transduced shRNA and verified its depletion in both J-Lat A2 and J-Lat 11.1 cells at the protein level by Western blotting (Figure 3A). Depletion of CHD9 led to a significant reversal of latency, as shown by an increase in the percentages of GFP positive cells (Figure 3B and C). Latency reversal was also confirmed by increased expression of viral genes Gag, Pol, and Tat in J-Lat 11.1 (Figure 3D). To characterize a potential direct association of CHD9 with the latent HIV-1 5’LTR, we performed chromatin immuno-precipitation (ChIP) in latent and PMA-activated 11.1 J-Lat cells. The positions of nucleosomes within the latent HIV-1 LTR are rigid, with positioned nucleosomes Nuc-0, Nuc-1, and Nuc-2 separated by DNAse I-sensitive regions HSS1 and HSS2, respectively, visually summarized in Figure 3E. CHD9 was found to be enriched throughout the HIV-1 LTR in latent 11.1 J-Lat cells, predominantly present over the Nuc0-HSS1 region, and this association was significantly decreased after LTR activation by PMA treatment (Figure 3F and Supplemental figure S5). No significant enrichment was observed at the HIV-1 internal control region (the GFP reporter) or the unrelated gene locus (HK2) (Figure 3F and Supplemental figure S5). As expected, PMA treatment led to an increase in histone H3 acetylation, a mark of active chromatin, over the HIV-1 5’LTR (Figure 3G and Supplemental figure S5). Our data demonstrate a role for CHD9 in the repression of HIV-1 gene expression and maintenance of latency via direct recruitment to and association with the HIV 5’LTR.

**Figure 3.**
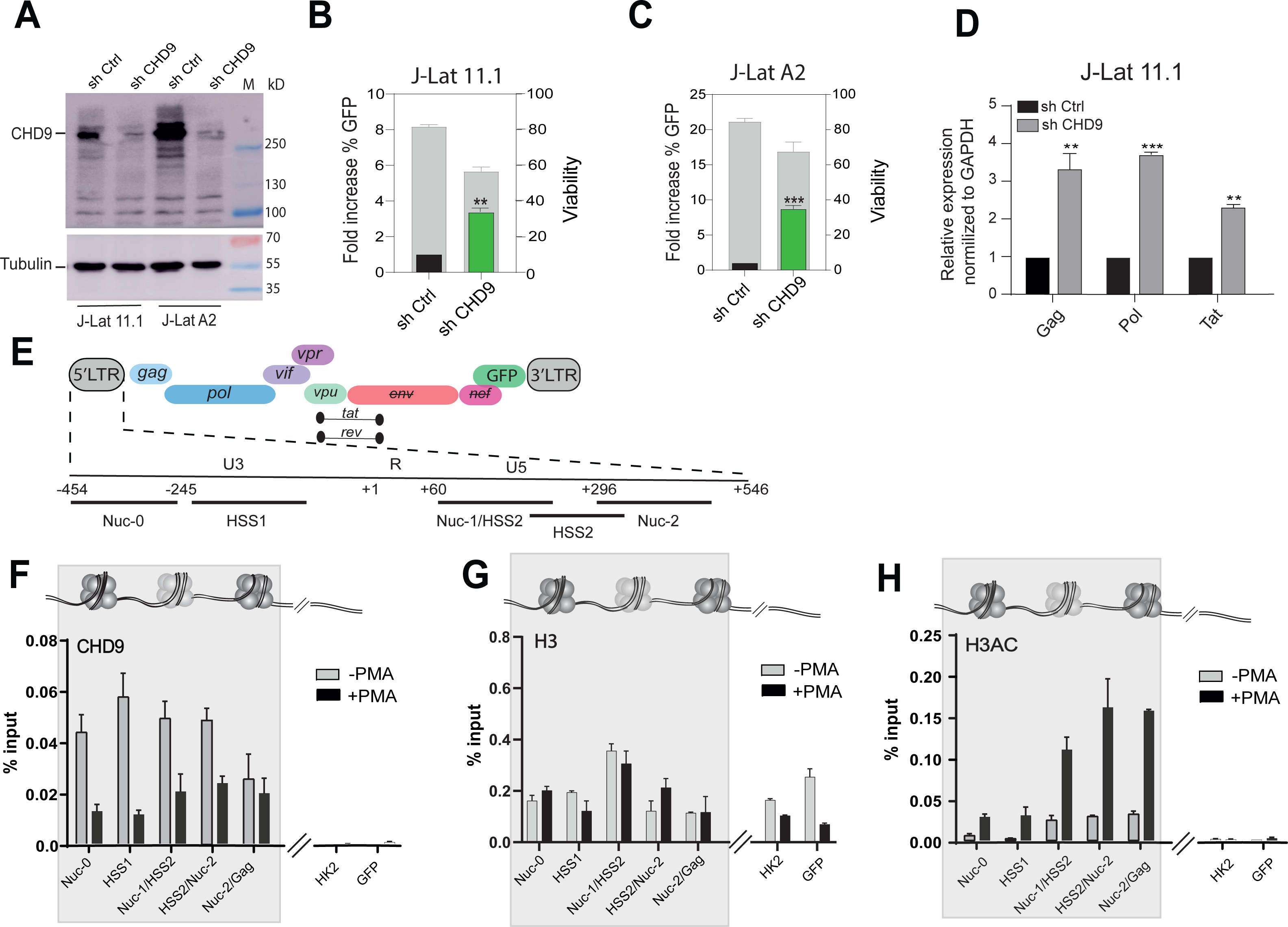
CHD9 regulates HIV-1latency in J-Lat A2s and J-Lat 11.1 cells. (A) Western blot for CHD9 in control and CHD9 shRNA depleted J-Lat 11.1 and A2 cells, α-Tubulin as a loading control. (B) Flow-cytometry bar plots demonstrating latency reversal after CHD9 shRNA depletion in J-Lat 11.1 cells: green bars indicate the percent of GFP positive cells after knockdown over control (black bar), left y-axis, whereas gray bars show cell viability, right y-axis. (C) Latency reversal after CHD9 shRNA depletion in J-Lat A2 cells. (D) Viral reactivation is confirmed by RT-qPCR in J-Lat 11.1 cells for the viral genes Tat, Gag and Pol. Data are normalized to GAPDH and represented as fold increase over sh Control. Statistical significance was calculated using ratio-paired t-test and multiple comparison t-test on Log2 transformed fold changes * – p < 0,05, ** – p < 0,01, *** – p < 0,001. (E) Schematic of HIV genome. 5’ LTR region further segmented into the U3, R, and U5 regions. Amplicons used in ChIP-qPCR experiments are indicated. (F) ChIP-qPCR analysis of CHD9 binding to the HIV-1 5’LTR in untreated and PMA stimulated J-Lat 11.1 cells. Data are represented as percentage of the input. (G) ChIP-qPCR analysis of Histone H3 occupancy at the HIV-1 5’LTR untreated and PMA stimulated J-Lat 11.1 cells. Data are represented as percentage of the input. (H) ChIP-qPCR analysis of Histone H3 acetylation at the HIV-1 5’LTR in untreated and PMA stimulated J-Lat 11.1cells. (E-H) Data represent the average (±SD) of two technical replicates.

### Pharmacological targeting of ADK, GRIK5 and NF1

With the aim of identifying potential latency reversing agents (LRAs), we performed a literature search and identified three candidate genes, ADK, GRIK5, and NF1, all present in the GFP^Sub^ list, for which FDA-approved small molecule inhibitors are available. We therefore examined the latency reversal potential of the ADK inhibitor 5-Iodotubercidin, the GRIK5 inhibitor Topiramate, and the NF1 inhibitor Trametinib. Adenosine kinase (ADK) is a phosphotransferase that converts adenosine into 5’-adenosine-monophosphate and thus plays a major role in regulating the intracellular and extracellular concentrations of adenosine, activation of specific signaling pathways, and bioenergetic and epigenetic functions (71, 72). 5-Iodotubercidin is a purine derivative that inhibits adenosine kinase by competing with adenosine for binding to the enzyme (73). Glutamate ionotropic receptor kainate type subunit 5 (GRIK5) is a subunit of the tetrameric kainate receptor (KAR), a subgroup of ionotropic glutamate receptors. GRIK5, together with GRIK4, binds glutamate, whereas subunits GRIK1-3 form functional ion-channels (74). Topiramate is an FDA-approved GRIK5 inhibitor employed as an anti-epileptic drug and is used to manage seizures and prevent migraines (75). Neurofibromin 1(NF1) is ubiquitously expressed; however, its highest levels are found in cells of the central nervous system and it has been described to function as a negative regulator of the Ras signal transduction pathway (76, 77). Trametinib is an NF1 inhibitor which is also known as a mitogen-activated protein kinase (MAPK) kinase (MEK) inhibitor with anticancer activity and is FDA-approved for use in metastatic malignant melanoma (78).

We examined the effects of treatment with 5-Iodotubercidin, Topiramate, and Trametinib in J-Lat 11.1, J-Lat A2 cells, and a primary CD4+ T cell model of latency, on latency reversal. Treatment with the ADK inhibitor 5-Iodotubercidin resulted in only a moderate induction of latency reversal in both A2 and 11.1 J-Lat cells and was accompanied by significant toxicity, especially at high concentrations [16μM] (Figure 4A & B). Only at a 5-Iodotubercidin concentration of 4μM in 11.1 J-Lat cells was a significant increase in GFP positive cells with acceptable viability observed (Figure 4A & B). Treatment of J-Lat A2 and 11.1 cells with the GRIK5 inhibitor Topiramate or the NF1 inhibitor Trametinib resulted in significant, concentration-dependent latency reversal (Figure 4A & B). Importantly, neither Topiramate nor Trametinib displayed significant toxicity in comparison with the DMSO vehicle control group, as measured by gating for live cells. In particular, treatment of J-Lat 11.1 cells with Trametinib resulted in significant latency reversal at all evaluated concentrations [250nM, 1μM, 4μM and 16μM], with minimal associated toxicity (Figure 4B).

**Figure 4.**
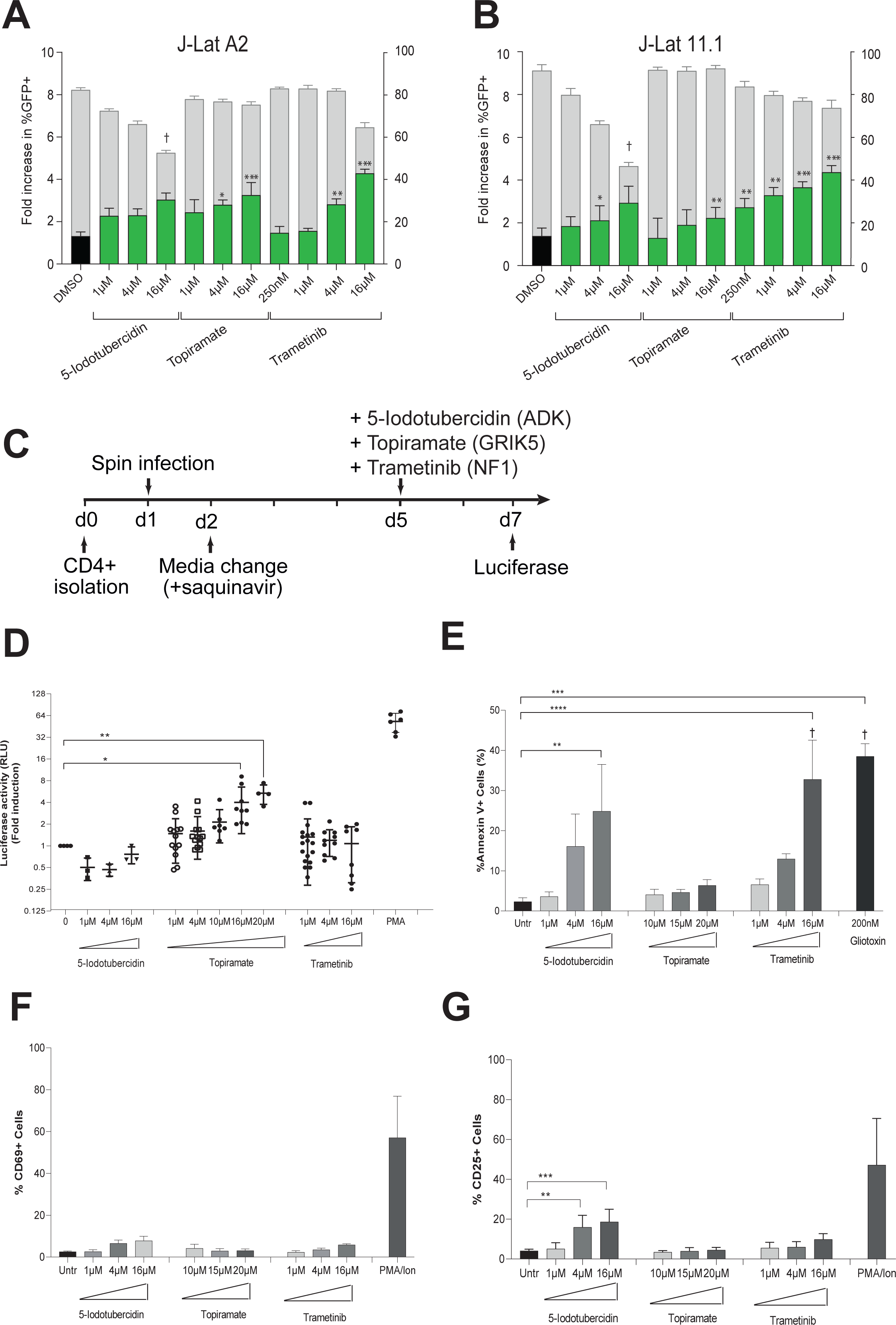
HIV latency reversal by small molecule inhibitors of three candidate genes ADK, GRIK5, NF1. (A) Latency reversal potential upon 48 h treatment of J-Lat A2 cells with increasing concentrations of 5-iodotubercidin (ADK inhibitor), Topiramate (GRIK5 inhibitor) and Trametinib (NF1 inhibitor) was evaluated by flow-cytometry. Treatment with DMSO (black bar) is used as a negative control. Percent of GFP positive cells is indicated by green bars (left y-axes), cell viability is indicated by grey bars (right y-axis). (B) Latency reversal potential upon 48 h treatment of J-Lat 11.1 cells. (C) Schematic representation of candidate LRAs treatment in a primary cell model of latent HIV-1 infection. CD4+ T-cells are isolated on day 0 and spin infected on day 1. On day 2 virus is removed by media change in presence of saquinavir. Latently infected cells are stimulated with candidate LRAs on day 5 and HIV reactivation is evaluated at day 7. (D) Latency reversal as measured by Luciferase activity in a primary cell model of HIV latency after treatment with 5-Iodotubercidin, Trametinib and Topiramate in different concentrations. Plots show the fold increase in luciferase activity, measured in relative light units (RLU), after treatment with different concentrations of 5-iodotubercidin (ADK inhibitor), Topiramate (GRIK5 inhibitor) and Trametinib (NF1 inhibitor). Each dot represents a single measurement, black horizontal lines show the average fold increase for each treatment. Averaged data of at least 3 independent experiments performed using each time two different donors (totaling at least 6 different donors). PMA was used as a positive control. Statistical significance was calculated using t test, * – p<0,05; ** – p<0,005; *** – p<0,0005. (E) Percentage of cells expressing apoptosis marker Annexin V in primary CD4+ T cells upon treatment with candidate LRAs for 48 hours. Treatment with a toxic concentration of Gliotoxin (GTX) 200nM was used as a positive control. Experiments were performed in uninfected cells obtained from 6 healthy donors. Data are presented as mean ±SD from three independent experiments. The (†) symbol indicates low viability. (F) Percentage of cells expressing marker of cell activation CD69 in primary CD4+ T cells from 6 healthy donors, data are presented as mean ±SD from three independent experiments of 2 different healthy donors upon treatment with candidate LRAs for 48 hours. Treatment with PMA/Ionomycin PMA is used as a positive control. (G) Percentage of cells expressing marker of cell activation CD25 in primary CD4+ T cells from 6 healthy donors, data are presented as Mean ±SD from three independent experiments of 2 different healthy donors upon treatment with candidate LRAs for 48 hours. Treatment with PMA/Ionomycin is used as a positive control. Statistical significance was calculated using one-way Anova, multiple comparison test. Asterisks indicate the level of significance. (**p < 0.01, ***p < 0.001).

Next, we evaluated the latency reversal potential of the inhibitors in a more clinically relevant primary ex vivo infection latency model, in which CD4+ T cells are infected with a full-length non-replication competent HIV-1 virus driving expression of a luciferase reporter (Figure 4C) (79). After spin infection, cells are allowed to rest for 4 days and are treated with the different compounds, as indicated, for 48 hours, followed by measurement of HIV-1 LTR-driven luciferase activity and staining for Annexin positive cells and T-cell activation markers CD69 and CD25 (Figure 4D-G and Supplemental figure S6). Similar to what was observed for the A2 and 11.1 J-Lat cell-lines, treatment of HIV-1-infected latent primary CD4+ T cells with the ADK-inhibitor 5-Iodotubercidin did not result in significant latency reversal (Figure 4D), while it produced significant toxicity at the highest concentration used, as indicated by Annexin V staining (Figure 4E). Treatment with the GRIK5 inhibitor Topiramate, at a concentration of 16 and 20 µM, resulted in significant (p < 0.05) reversal of latency, as measured by a 3-6 fold increases in mean luciferase activity compared to untreated controls (Figure 4D). Topiramate treatment did not significantly affect CD4+ T cell viability after 48 hours, while control treatment with a toxic (200nM) concentration of Gliotoxin (GTX) caused apoptosis of primary CD4+ T-cells, as evidenced by an increase in the percentage of Annexin V-positive cells, as was observed previously (Figure 4E) (80). In contrast to what was observed in the A2 and 11.1 J-Lat cell-lines, treatment of latently infected CD4+ T-cells with the NF1 inhibitor, Trametinib did not lead to a significant reversal of latency (Figure 4D), while viability was only moderately affected (figure 4E).

For LRA candidates to be viable in a clinical setting, it is crucial that they reactivate latent HIV without inducing activation of CD4+ T-cells. Therefore, we examined the potential of our candidate LRAs to induce expression of the activation markers CD25 and CD69 in treated CD4+T cells. While the ADK inhibitor 5-Iodotubercidin resulted in markedly elevated expression of the activation markers, the GRIK5 inhibitor Topiramate and NF1 inhibitor Trametinib did not induce significant induction of the T cell activation markers CD69 and CD25 (Figure 4F & G and Supplemental figure S6). As expected, PMA/Ionomycin treatment substantially activated T cells (Figure 4F & G and Supplemental figure S6). Taken together, our data indicate that the FDA-approved GRIK5 inhibitor Topiramate reverses HIV-1 latency in a variety of T cell models of latency, without induction of T cell activation and with limited cytotoxicity, and, therefore, can be considered an attractive LRA for further mechanistic and pre-clinical investigation.

## Discussion

In search of potentially novel host factors and pathways that play a role in the maintenance of HIV-1 latency, we performed a two-color haploid genetic screen in latent HIV-1-infected KBM7 cells. An important advantage presented by this approach is that identification of putative functionally relevant candidate latency-promoting host target genes does not require a-priori knowledge of the molecular determinants of latency and is thus completely unbiased. Additionally, gene-trap insertional mutagenesis has enabled identification of previously unappreciated latency genes and cellular pathways, which likely modulate latency not only via direct physical association with the HIV-1 promoter, but also indirectly, through involvement in cellular signaling. We produced a list of 69 candidate genes and proceeded to validate ten candidates, the depletion of which in various in vitro models of latency, including primary CD4+ T-cell models, led to latency reversal.

The haploid KBM7 cell line, while a powerful system for GT-mediated forward genetic screens, is a myeloid cell line (36). HIV can infect and establish latent infection in monocytes and macrophages (81, 82), but these are not considered to be the prime source of the latent reservoir; this begs the question how relevant the myeloid nuclear environment may be for HIV latency in lymphocytic cells. Nevertheless, when we tested a selected subset of the candidate genes by shRNA-mediated knockdown in a T-cell derived model of HIV latency, we found that knockdown of 67% of candidates (10 out of 15) led to latency reversal, which demonstrates the validity of our approach. Moreover, several of the genes identified in our gene trap screen have previously been implicated in HIV susceptibility: 15 out of 69 genes are listed in the HIV-1 human interaction database (https://www.ncbi.nlm.nih.gov/genome/viruses/retroviruses/hiv-1/interactions/), a detailed database of all known interactions between HIV-1 and the human host. Furthermore, one of the candidates, IRF2BP2, is a potential target of HIV-associated nucleotide polymorphisms within a cluster of regulatory DNA elements (66) which loop to and potentially regulate the IRF2BP2 promoter in CD4+ T-cells (83, 84). In the current study, we demonstrate that knock-down of IRF2BP2 results in latency reversal. IRF2BP2 has been shown to interact with NFAT1 and to repress transcriptional activity (85), providing a plausible mechanism for its role in latency maintenance.

Among the candidate gene list, we also identified CHD9, a member of the chromodomain helicase DNA-binding (CHD) family of the ATP dependent chromatin remodelers. Members of this family are involved in various cellular processes and in normal development and disease; however, CHD9 is one of the least-studied members. We found that CHD9 is associated with the latent HIV-1 5’LTR and is displaced upon promoter activation by PMA stimulation, suggesting that it acts as a repressor of HIV transcription. Indeed, depletion of CHD9 by shRNA-mediated knockdown led to de-repression of HIV, as observed by increase in the expression of the HIV-1 LTR-driven reporter GFP as well as the HIV-1 genes Gag, Pol and Tat. Future studies will determine the mode of recruitment of CHD9 to the HIV-1 LTR and the molecular interplay between CHD9 and other chromatin remodeler and modifying complexes associated with the latent HIV-1 5’LTR (4).

A large subset of the list of 69 candidate genes are non-coding RNAs (N=22); upon closer inspection we found that 16 of those are at least in part oriented in an anti-sense direction with respect to known protein-coding genes. We currently do not know if these non-coding transcripts fulfill a biological function in maintenance of HIV-1 latency, directly through their transcripts, through regulatory effects on other (protein-coding) genes in cis or in trans, through other effects, or if they represent mapping artefacts. Further investigation into these possibilities is ongoing.

An interesting observation emerging from our experimental set-up is the appearance of a sub-population of cells after multiple rounds of sorting that is more stable in its GFP expression. This sub-population is also higher in mCherry expression, as compared to the total double-positive population. The reversal of double-positive cells after sorting may reflect the intrinsically stochastic transcription of HIV-1(86–88). Latent cell lines are notoriously sensitive to cellular stresses, which cause reactivation (89–91). It is, therefore, possible that the less stable GFP-positive cells represent cells that are temporarily activated, and which slowly revert back to a latent state. Therefore, we focused our analysis and validation experiments mainly on the stable GFP-high population. Nevertheless, the few candidate genes unique to the total population that we tested in our shRNA knockdown validation experiment had a similar false positive ratio (i.e. 33.3%) as we found for the candidate genes from the stable GFP-positive population.

For our analysis, we set a strict criterion of the LOF score being ≥3 in at least two biological replicates, although a cut-off at lower LOF scores of 2 or above has been used previously (92). We chose this stringent criteria to confidently identify 69 candidate genes. We believe however that our dataset likely contains additional candidate genes that we miss due to the stringency applied. This is exemplified by our observation that STRING analysis indicates that many of the 598 protein-coding genes with an LOF score of 3 and higher functionally interact (Supplemental Figure S7). Moreover, 32% of these genes (195) appear in the HIV-1 human interaction database, pointing to their potential roles in HIV-1 biology. Importantly, proteins with well-described functions in HIV biology, such as, e.g., IL32 (LOF=9.5; (66), RUNX1 (LOF=12.6; (93), and SMARCA4/BRG1 (LOF=8.7; (59), are detected with high LOF scores in at least one of our samples.

Identification of functionally relevant molecular targets and novel compounds effective in latency reversal has proven to be an outstanding challenge in the field (94). Most clinically investigated LRAs have thus far failed to significantly impact the size of the latent reservoir in patients. From our list of validated host factors we identified three druggable targets, ADK, NF1, and GRIK5. The ADK inhibitor 5-Iodotubercidin displayed cytotoxicity in both J-Lat and primary models of HIV latency. Since knockdown of ADK did not lead to significant toxicity, off target, pleiotropic effects of 5-Iodotubercidin may underlie the observed T cell toxicity. 5-Iodotubercidin is the prototype nucleoside ADK inhibitor, with a relatively high IC_50_. Novel nucleoside and non-nucleoside ADK inhibitors have been developed with more favorable pharmacological properties that would be attractive to pursue in the context of HIV-1 latency reversal in future studies (72). Trametinib, a small molecule inhibitor of NF1, reactivated HIV-1 in J-Lat cell models with modest effects on viability but displayed significant toxicity in the primary cell models. These results highlight the importance of interrogating the potential effectiveness of candidate LRAs in multiple models of HIV-1 latency to circumvent potential integration-site mediated biases present in cell line models of latency, and, most importantly, to confirm effectiveness in the more relevant primary CD4+ T cells harboring latent HIV-1 in vivo.

Our data point to the GRIK5 inhibitor Topiramate as a potentially promising compound for latency reversal. Topiramate, effectively reversed latency in primary HIV-1-infected CD4+ T cells without inducing significant T cell activation or cytotoxicity; this makes Topiramate a potentially clinically promising LRA and a target for further investigation. GRIK5 (glutamate ionotropic receptor kainate type subunit 5), primarily studied in neurons, is a subunit of the tetrameric kainate receptor (KAR), a subgroup of ionotropic glutamate receptors. GRIK5, together with GRIK4, bind glutamate, whereas subunits GRIK1-3 form functional ion-channels (74). In B-cells, KAR activation by glutamate increases ADAM10 levels, leading to increased B cell proliferation and immunoglobulin production (95). Topiramate is primarily used as an anticonvulsant or antiepileptic drug. While the exact mechanism by which Topiramate exerts anticonvulsant or antiepileptic properties is unclear, it has been shown to block voltage-dependent sodium and calcium channels (73, 96), to inhibit the excitatory glutamate pathway and enhance inhibition by GABA (97). Interestingly, Topiramate can induce cytochrome P450 family member CYP3A4 activity and potentially negatively affect the metabolism of many drugs (98), which should be taken into account when considering a potential therapeutic combination of LRAs in future studies.

## Acknowledgments

TM received funding from the European Research Council (ERC) under the European Union’s Seventh Framework Programme (FP/2007-2013)/ERC STG 337116 Trxn-PURGE, Dutch AIDS Fonds grant 2014021, and Erasmus MC mRACE research grant. RJP received funding from Dutch AIDS Fonds grant 2016014. We thank Thijn Brummelkamp for providing materials and expertise.

## Author Contributions

MR, MMS, EN, MS, EDC, HB,TWK, MA, RJP and TM carried out experiments and performed data analysis. PM, VH, and PH performed Nextgen sequencing and data analysis. MR, EN, MMS, RJP and TM conceived the study and wrote the manuscript. All authors read and approved the final manuscript.

## Materials and Methods

### Cell culture

KBM7 (a kind gift from Thijn Brummelkamp) and Hap-Lat (latent HIV infected KBM7-derived cell lines) were cultured in IMDM media (ThermoFisher Scientifc) supplemented with 10% FCS and 2% Pen/Strep. Haploidy of KBM7 cells was maintained by periodically sorting cells for size (5% smallest). Ploidy of cells was determined by propidium iodide staining

### Establishment of haploid latent (Hap-Lat) HIV infected cell lines

We used a strategy described previously (58, 59) to generate latently HIV infected KBM7 haploid cell lines. Minimal HIV (LTR-GFP, HIV-658 (99) virus was produced by transfection of 293T cells in 15cm culture dishes using polyethylenimine (PEI) Sigma Aldrich) with a mixture of 6.8μg p658, 2μg VSVG and 4.5μg Gag-pol plasmids. Haploid KBM7 cells were infected with HIV-derived virus at low MOI such that approximately 5-10% of cells became productively infected as determined by GFP expression. 5 days after infection, the GFP negative cell population harboring uninfected and potentially latently infected cells were sorted by flow cytometry activated cell sorting (FACS) and stimulated with 350nM SAHA (Selleck Chem), and 5ng/µl TNFα (Sigma-Aldrich). Twenty-four hours post stimulation, GFP positive cells were single cell sorted by FACS into 96 well plates. Clones were expanded and characterized for their basal GFP expression and their potential for re-activation. From the clonal cell lines generated Hap-Lat #1 was selected for low background and relative high reactivation upon stimulation (GFP negative under basal conditions but inducible to express GFP upon activation).

### Gene Trap virus production and mutagenesis of Hap-Lat cell lines

We adapted a strategy described previously (37) to mutagenize Hap-Lat cells. Briefly, Gene Trap virus was produced by transfection of HEK293T cells in 15cm culture dishes using 180 μl polyethylenimine (PEI) with a mixture of 8 μg pGT-mCherry, 2.1 μg pAdvantage, 3.1 μg CMV-VSVG and 4.8 μg Gag-pol plasmids in at total volume of 1ml serum free RPMI. After 12 hours the growth media was changed to fresh FCS supplemented RPMI. The virus was collected at 12, 24, 36 and 48 hours. For the GT mutagenesis, 192 million Hap-Lat #1 cells were pre-incubated with 100μM dNTPs (Invitrogen) 1hr before transduction. Cells are resuspended in 192ml of undiluted GT virus. Spin infection was performed for 90minutes @ 1500rpm in 24 well plates containing 2 million cells per well and 8μl protamine sulfate per ml. After 24 hours cells were subjected to a second round of infection after which virus was removed, and cells were left to recover in supplemented IMDM. We estimate that this sequential infection resulted in 1 to 4 GT integrations per cell, with the majority of cells containing one integration (data not shown). A first round of sorting was performed 8 days after the second infection. Subsequently, between 500.000 and 1 million GFP-mCherry double positive cells were sorted each day for 5 consecutive days using two cell sorters (BD FACSAriaII SORP and BD FACSAriaIII). Sorted cells were pooled, left to recover and expanded for the next round of sorting. A second round of sorting was performed 17 days after the second infection. Between 2.6 and 3 million cells were collected each day for 3 consecutive days using two sorters. These cells were pooled and left to recover and expand for a third round of sorting on day 22 after the second infection, yielding 6.1 million cells from 1-day sorting on two machines. On day 29 after the second infection a fourth round of sorting was performed, yielding 7.2 million cells from 1-day sorting on two machines. After the cells of the fourth round of sorting were left to recover and expand a sub-population of highly mCherry and GFP double positive cells were apparent. In a final fifth round of sorting the total GFP-mCherry positive population (10.1 million cells) and the GFP-mCherry positive high sub-population (5.9 million cells) were respectively sorted on day 33 and 35 after the second infection. After recovery, genomic DNA was isolated from the sorted cell populations, as well as from a population of mutagenized unsorted cells.

### Mapping insertion sites through inverse PCR

A previously described inverse PCR protocol was adapted to determine host sequences flanking the proviral insertion sites (37). Briefly, genomic DNA was isolated from 5 million cells using the DNAeasy kit (Qiagen). 4ug gDNA was digested with NlaIII or MseI. After PCR spin column purification (Qiagen), 50ul of eluted digested DNA was ligated using T4 DNA ligase (Roche) in a volume of 2ml. The reaction mix was purified using spin columns and used in as template in a PCR reaction with primers annealing to internal sequences in the gene trap vector (5’-CTGCAGCATCGTTCTGTGTT-3’ and 5’-TCTCCAAATCTCGGTGGAAC-3’). The resulting PCR products were used for library preparation.

### High-throughput sequencing and identification of integration sites

Sequencing libraries were created using the Ion Plus Fragment Library Kit (ThermoFisher Scientific) according to manufacturer’s instructions, with minor modifications: briefly, 15ng of the PCR products were diluted with ddH2O to a final volume of 39 μl. The samples were end-repaired, adaptor ligated and amplified; this was followed by 2 rounds of purification using Agencourt AMPure XP beads. Library qualities and quantities were assessed by Bioanalyzer, using the DNA High Sensitivity Kit (Agilent Technologies). The quantified libraries were pooled together in 10plex, at a final concentration of 40pM. Templating, enrichment and chip loading was performed on an Ion Proton Chef system using the Ion PI™ Hi-Q™ Chef Kits (ThermoFisher Scientific); sequencing was performed on an Ion Proton PI™ V3 chip, with the Ion PI™ Hi-Q™ Sequencing 200 Kit, on an Ion Proton instrument (ThermoFisher Scientific), according to manufacturer’s instructions.

The resulting FASTQ files were processed to remove the gene trap vector primers. To this end, we used R/Bioconductor packages ShortRead (100) and Biostrings to match primer sequences within the sequenced reads, allowing for two mismatches at maximum for each primer sequence. The identified primers were trimmed and reads shorter than 15bp were discarded. The remaining reads were aligned to the human reference genome version hg19 retrieved from the Illumina iGenomes repository, using bwa mem. The produced aligned reads (BAM files) were then subjected to integration site identification. To this end, we re-implemented the HaSAPPy algorithm (64) in R. The re-implemented version operates on BAM instead of SAM files and was enriched with additional noise filtering steps, which improved the overall process of detecting integration sites using the Local Outlier Method (LOF), as described in the original HaSAPPy implementation. Specifically, we added attributes for i) filtering out read artifacts from introns, based on the overall intronic read distribution for each sample, ii) selecting the fraction of neighboring (Independent Insertions, Disrupting Insertions, Bias) triplets for LOF analysis and iii) library normalization across samples. After the quality control steps, our HaSAPPY re-implementation assigns a Local Outlier Factor (LOF) score to each gene in a sample based on a triplet score derived from the number of putative GT integrations in the sample compared to the reference, additionally taking into account the potential bias arising from unbalanced sense and anti-sense read numbers. LOF is a metric to indicate to what extent a vector measurement deviates from a population of vectors with the same properties. In our case, a triplet vector corresponds to a gene and LOF measures the deviation from the respective population, i.e. a gene is an outlier based on a distribution of vectorized scores, such as the aforementioned triplet. The re-implemented version can be found in https://github.com/pmoulos/ngs-stone-age/blob/master/R/hasar.R.

### ShRNA knock-down of candidate genes in A2 and 11.1 cell lines

Pre-designed shRNA sequences (MISSION® shRNA library (Sigma); Table 2) were purchased as bacterial glycerol stocks from the Erasmus Center for Biomics. Virus was produced as follows, 3 × 10^5^ HEK293T cells/ml were plated in a 10 cm dish cells one day before transfection and transfected with 4,5 μg of pCMVΔR8.9 (envelope), 2 μg of pCMV-VSV-G (packaging) and 6ug of shRNA vector. The transfection mixture was removed after 12 hours and replaced with fresh RPMI medium contain 10% FBS. Virus containing supernatant was collected at 36, 48, 60- and 72-hours post-transfection. Jurkat, J-Lat A2 (LTR-Tat-IRES-GFP) and J-Lat 11.1 (integrated full-length HIV-1 genome mutated in env gene and GFP replacing Nef) cells were cultured in RPMI-1640 medium (Sigma) supplemented with 10% FBS and 100 μg/ml penicillin-streptomycin at 37 °C in a humidified 95% air-5% CO2 atmosphere. A2 and 11.1 cell lines were infected by adding 500 µl of filtered lentivirus to 2.5 ml of cell culture. After 48 hours, medium was refreshed and cells were puromycin selected. 14 days after infection, RNA was harvested to determine knock-down of genes by qPCR and reactivation of latent HIV was measured by flow-cytometry and RT-qPCR for Gag, Pol and Tat.

**Table 2.** List of Mission shRNA clones.

| **Recombinant DNA** |  |  |
| --- | --- | --- |
| sh ADK | MISSION® shRNA library (Sigma) | TRCN000011535 |
| sh GRIK5 | MISSION® shRNA library (Sigma) | TRCN000011537 |
| sh SPN | MISSION® shRNA library (Sigma) | TRCN000011539 |
| sh Control | MISSION® shRNA library (Sigma) | SHC002 |
| sh CMSS1 | MISSION® shRNA library (Sigma) | TRCN000011541 |
| sh EVL | MISSION® shRNA library (Sigma) | TRCN000011542 |
| sh COPS5 | MISSION® shRNA library (Sigma) | TRCN000011543 |
| sh EXOSC8 | MISSION® shRNA library (Sigma) | TRCN000011544 |
| sh SLC7A6 | MISSION® shRNA library (Sigma) | TRCN000011545 |
| sh USP15 | MISSION® shRNA library (Sigma) | TRCN000011547 |
| sh NF1 | MISSION® shRNA library (Sigma) | TRCN000011638 |
| sh IRF2BP2 | MISSION® shRNA library (Sigma) | TRCN000011635 |
| sh RHOF | MISSION® shRNA library (Sigma) | TRCN000011642 |
| sh EVI2B | MISSION® shRNA library (Sigma) | TRCN000011644 |
| sh CHD9 | MISSION® shRNA library (Sigma) | TRCN0000230236 |
| sh FAM19A | MISSION® shRNA library (Sigma) | TRCN000011641 |

### Flow cytometry for GFP expression in the J-Lat cell lines

Cells were collected in PBS. GFP fluorescent signal indicating latency reversal was monitored using a LSRFortessa (BD Biosciences). Viability was determined using the forward scatter area versus side scatter area profiles (FSC-A, SSC-A). Data was analyzed using FlowJo software (version 9.7.4, Tree Star).

### Total RNA Isolation and Quantitative RT-PCR (RT-qPCR)

Total RNA was isolated from transduced A2 and 11.1 cells using Trizol (Thermo Fisher) on day 14 after infection using Total RNA Zol out kit (A&A Biotechnology), residual genomic DNA was digested with DNAseI (Life technologies). cDNA was synthesized using superscript II Reverse Transcriptase (Life Technologies) using oligo (dT) primers or random primers (for Gag, Pol and Tat). RT-qPCR reactions were performed on a CFX Connect Real-Time PCR Detection System thermocycler (BioRad) using GoTaq qPCR Master Mix (Promega) (3 min at 95 °C, followed by 40 cycles of 95 °C for 10 s and 60 °C for 30 s). Melting curve analysis was performed to assess specificity of RT-qPCR products. Primers used for real-time PCR are listed in Table 3. Expression data was calculated using 2-ΔΔCt method (101). ß-2-microglobulin (B2M) and GAPDH were used as housekeeping genes for the analysis.

**Table 3.**
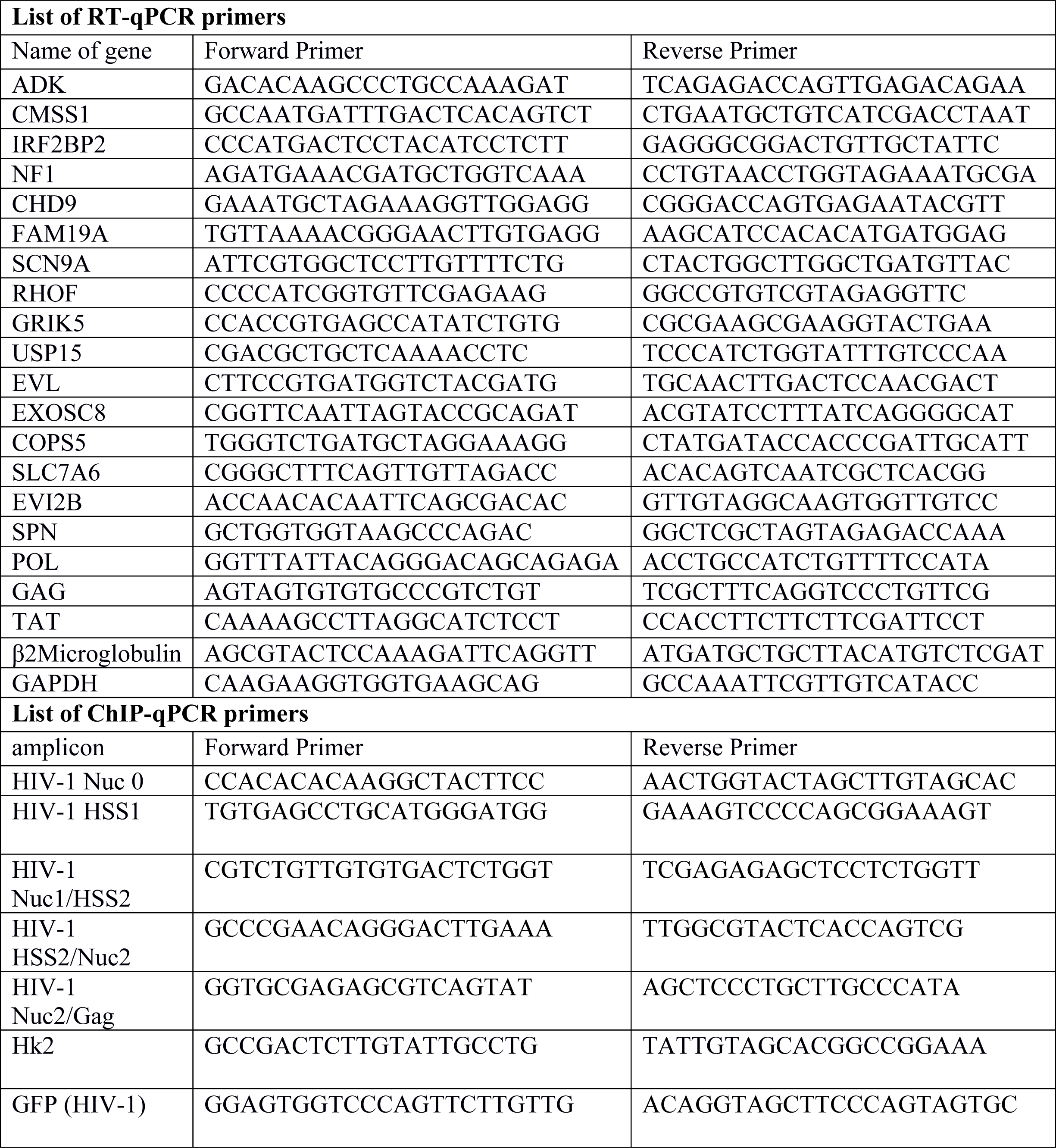
List of qPCR primers.

### Isolation and ex vivo infection of primary CD4+ T cells

HIV-1 latency *ex vivo* model was generated based on Lassen and Greene method by spinoculation (90 min at 1200 g), washed in PBS and cultured in RPMI 1640 containing 10% FBS, 100 μg/ml penicillin-streptomycin, IL2 (5ng/ml) and the antiretroviral drug Saquinavir Mesylate (5μM) (79). HEK 293T cells were transfected with HXB2 Env and pNL4.3.Luc.R-E-plasmids using PEI (Polyethylenimine) as described above. Supernatants containing pseudovirus particles are collected at 24, 48 and 72 h post-transfection. Peripheral blood mononuclear cells (PBMCs) were isolated by Ficoll density gradient sedimentation of buffy coats from healthy donors. Total CD4+ T cells were separated by negative selection using RosetteSep™ Human CD4+ T Cell Enrichment Cocktail (StemCell Technologies). Primary CD4+ T cells were plated at a concentration of 1,5×10 ^6^ cells/mL left overnight for recovery and infected. Two days after infection, cells were collected, washed once with PBS and treated with 5-Iodotubercidin, Topiramate or Trametinib. Cells were collected in 48 hr after treating, washed once in PBS and luciferase activity was measured using Luciferase Assay System (Promega).

### Flow cytometry for T cells activation and toxicity assay

Primary CD4+ T cells isolated from buffy coats of healthy volunteers were treated with different concentration of 5-Iodotubercidin, Topiramate and Trametinib and PMA/ionomycin as a positive control.

After 24 and 48 hours, cells were stained with Annexin V to examine the percentage of cells undergoing apoptosis and with the surface receptor CD69 and CD25 to measure T cell activation. For staining, 10^6^ cells were washed with PBS supplemented with 3% FBS and 2.5 mM CaCl2 followed by staining with Annexin V-PE (Becton and Dickinson), CD69-FITC (eBioscience) and CD25-APC (eBioscience) for 20 minutes at 4C in the presence of 2.5 mM CaCl2. Cells were then washed with PBS/FBS/CaCl2 and analyzed by flow cytometry. Between 2 - 4×10^5^ events were collected per sample within 3 hours after staining on a LSRFortessa (BD Biosciences, 4 lasers, 18 parameters) and analyzed using FlowJo software (version 9.7.4, Tree Star).

### Western Blotting

10×10^6^ cells were lysed for 30 minutes on ice with 200 ul lysis buffer (150 mM NaCl, 30 mM Tris (pH 7.5), 1 mM EDTA, 1% Triton X-100, 10% Glycerol, 0.5 mM DTT, protease inhibitor cocktail tablets (EDTA-free) (Roche). Cell lysates were clarified by centrifugation (14,000 rpm for 30 min at 4°C), mixed with 4x sodium dodecyl sulfate (SDS)-loading buffer containing 0.1M DTT and boiled at 95°C for 5 min. Samples were run in a 10% SDS-polyacrylamide gel at 100V. The proteins were transferred to polyvinylidene difluoride (PVDF) membranes and the membranes were probed with anti-CHD9 (13402-1-AP, Proteintech) and anti-α-tubulin (ab6160, Abcam). The next day, blots were incubated with HRP conjugated secondary antibody and proteins were imaged by chemical luminescence using Super Signal West Pico (Thermo Scientific).

### Chromatin immunoprecipitation (ChIP) and qPCR

Approximately 50 to 100 million J-Lat 11.1 cells per condition were collected and crosslinked with 1% formaldehyde (Polysciences inc.) in 40 ml PBS, supplemented with 1mM MgCl_2_ and 1mM CaCl_2_, at RT for 30 min with 15rpm vertical rotation. The reaction was quenched with 125 mM Glycine. Cross-linked cells were pelleted at 800rcf for 5 minutes RT and subjected to chromatin enrichment as described previously (102). Briefly: the pellets were re-suspended in 2 ml ChIP incubation buffer (1% SDS, 1% Triton x-100, 150mM NaCl, 1mM EDTA, ph 8.0, 0.5 mM EGTA, 20mM HEPES pH 7.6, protease inhibitor cocktail tablets (EDTA-free, Roche) containing 1% SDS and sonicated (30’ ON/30’ OFF intervals Diogenode Bioruptor plus) to obtain chromatin fragments between 300 and 500bp. Sonicated chromatin was spun at 20817 rcf at 4°C for 30 minutes and diluted 10 times in ChIP incubation buffer without SDS. Diluted chromatin corresponding to 30-40 million cells was precleared overnight at 4°C with vertical rotation at 15rpm with 80uls of Protein A SepharoseTM 4 Fast flow (GE Healthcare) beads and immunoprecipitated overnight with 40uls of beads and 5ug of antibody (13402-1-AP, Proteintech). Samples were washed two times per each buffer (5minutes, 15rpm vertical rotation) with buffer 1 (0.1% SDS, 0.1% DOC, 1% Triton x-100, 150mM NaCl, 1 mM EDTA pH 8.0, 0.5 mM EGTA, 20mM HEPES pH8.0.), buffer 2 (500mM NaCl: 0.1% SDS, 0.1% DOC, 1% Triton x-100, 500mM NaCl, 1 mM EDTA pH 8.0, 0.5mM EGTA, 20mM HEPES pH8.0), buffer 3 (0.25M LiCL, 0.5%DOC, 0.5% NP-40, 1mM EDTA pH 8.0, 0.5mM EGTA pH 8.0, 20mM HEPES pH 8.0), buffer 4 (1mM EDTA pH8.0, 0.5mM EGTA pH 8.0, 20mM HEPES pH 8.0) to remove unspecific binding. Finally, each sample was eluted in 400ul elution buffer (1% SDS, 0.1M NaHCO_3_) de-crosslinked overnight at 65°C in presence of 200mM NaCl, phenol-chloroform extracted, ethanol precipitated and subjected to qPCR analysis using the primer sets summarized in Table 3.

## Figure legends

**Supplemental Figure 1.**
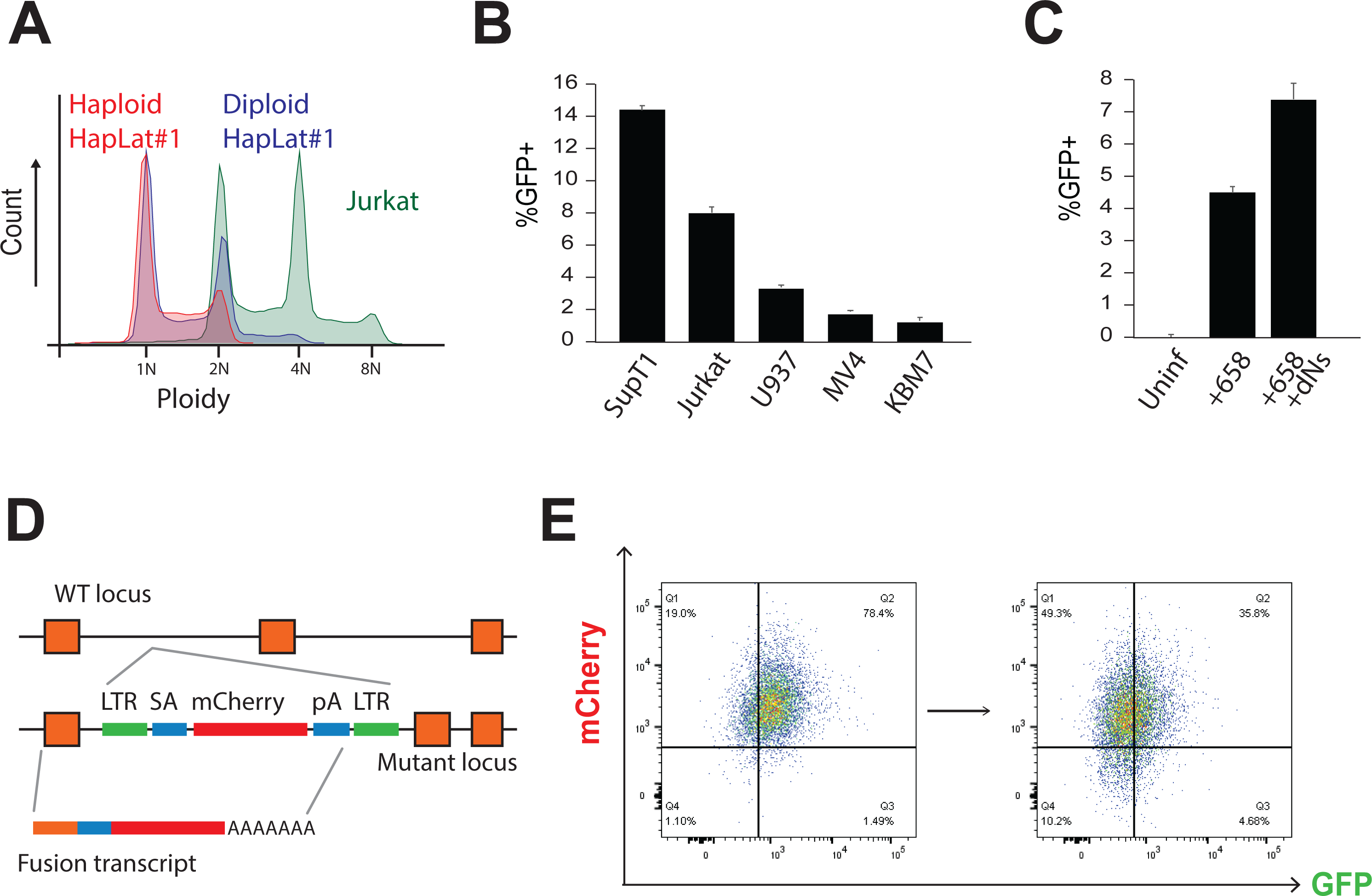
Generation of haploid latent cell lines. (A) Ploidy of Hap-Lat #1 compared to Hap-Lat #1 cells cultured over a prolonged period and Jurkat cells. Haploidy of HapLat #1 cells can be maintained over time by periodic cell sorting of small cells. (B) Infectivity to HIV-1 of KBM7 and other myeloid lineage cell lines, U937 and MV4, is poor compared to lymphocyte derived cell lines (SupT1 and Jurkat) as shown by percentage GFP positive cells after 4 days of infections with minimal HIV (658). (C) Nucleoside pre-incubation before and after supplementation during infection improves infectivity of KBM7 cells. (D) Schematic representing the mechanism of GT insertional mutagenesis. The inactive LTRs of MSCV flank a splice acceptor (SA) site, a mCherry reporter and a poly-adenylation (pA) terminator, (E) After sorting and expansion, sorted double-positive cells tend to revert to a latent state as indicated by a loss of GFP signal. FACS plots obtained 1day (left panel) and 5 days (right panel) after sorting.

**Supplemental Figure 2.**
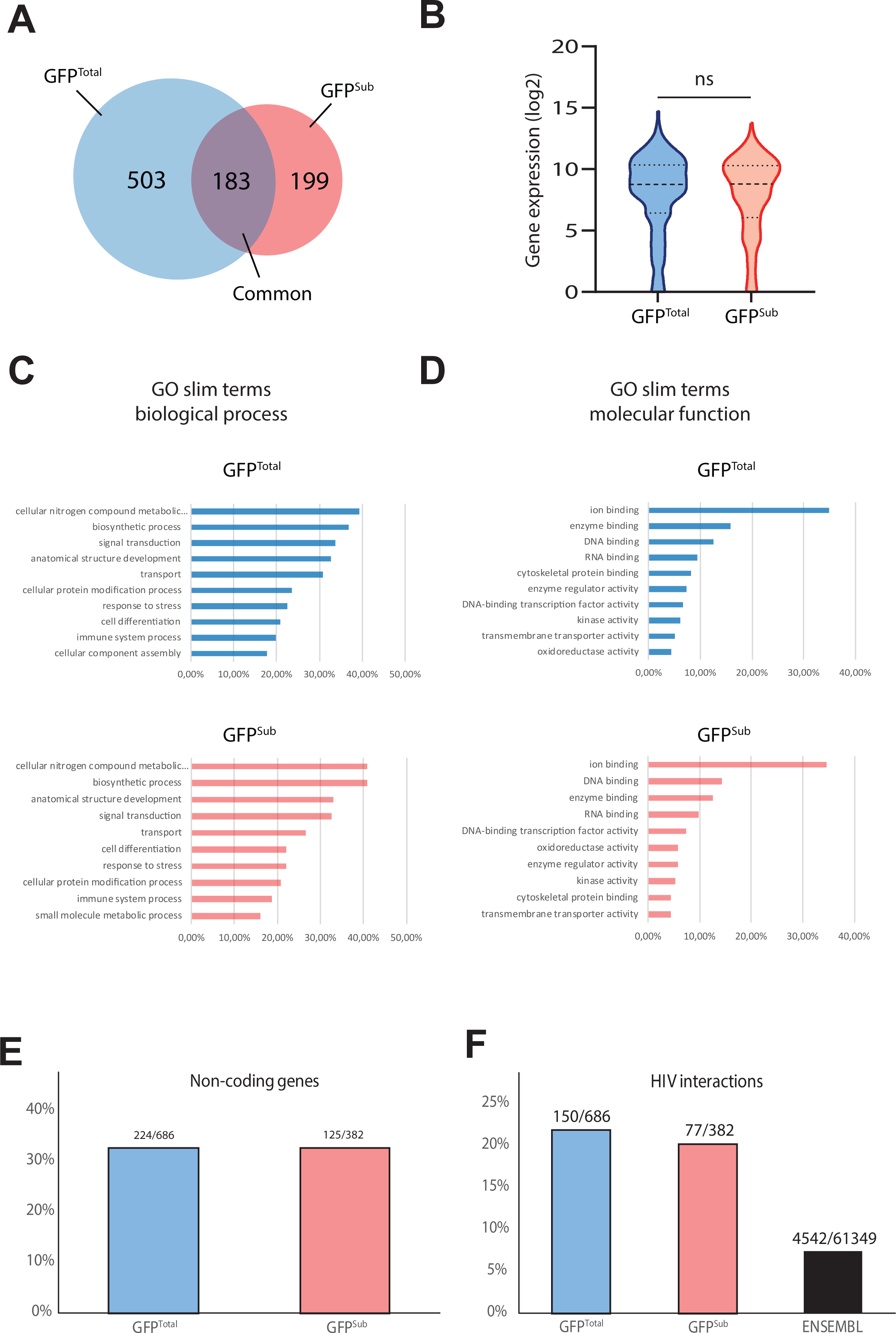
Characterization of gene trap integration candidate genes. (A) Venn diagram depicting all genes with an LOF score >3 in one replicate from the GFP^Total^ (686 hits) and the GFP^Sub^ (382 hits) population. 183 genes are common to both populations. (B) Expression levels of genes in the GFP^Total^ and the GFP^Sub^ populations. No significant differences in the average expression levels of genes between the two populations is found (Mann-Whitney test). (C) GO-slim term analysis for biological process of genes in the GFP^Total^ and the GFP^Sub^ populations. No substantial differences in GO-slim term enrichment are found. (D) GO-slim term analysis for molecular function of genes from both populations. (E) Percentage of non-coding genes present among genes in the GFP^Total^ and the GFP^Sub^ populations are similar. (F) Percentage of genes present in the HIV interaction database (https://www.ncbi.nlm.nih.gov/genome/viruses/retroviruses/hiv-1/interactions/) among genes in the GFP^Total^ and the GFP^Sub^ populations are similar but higher compared to the total ENSEMBL gene list.

**Supplemental Figure 3.**
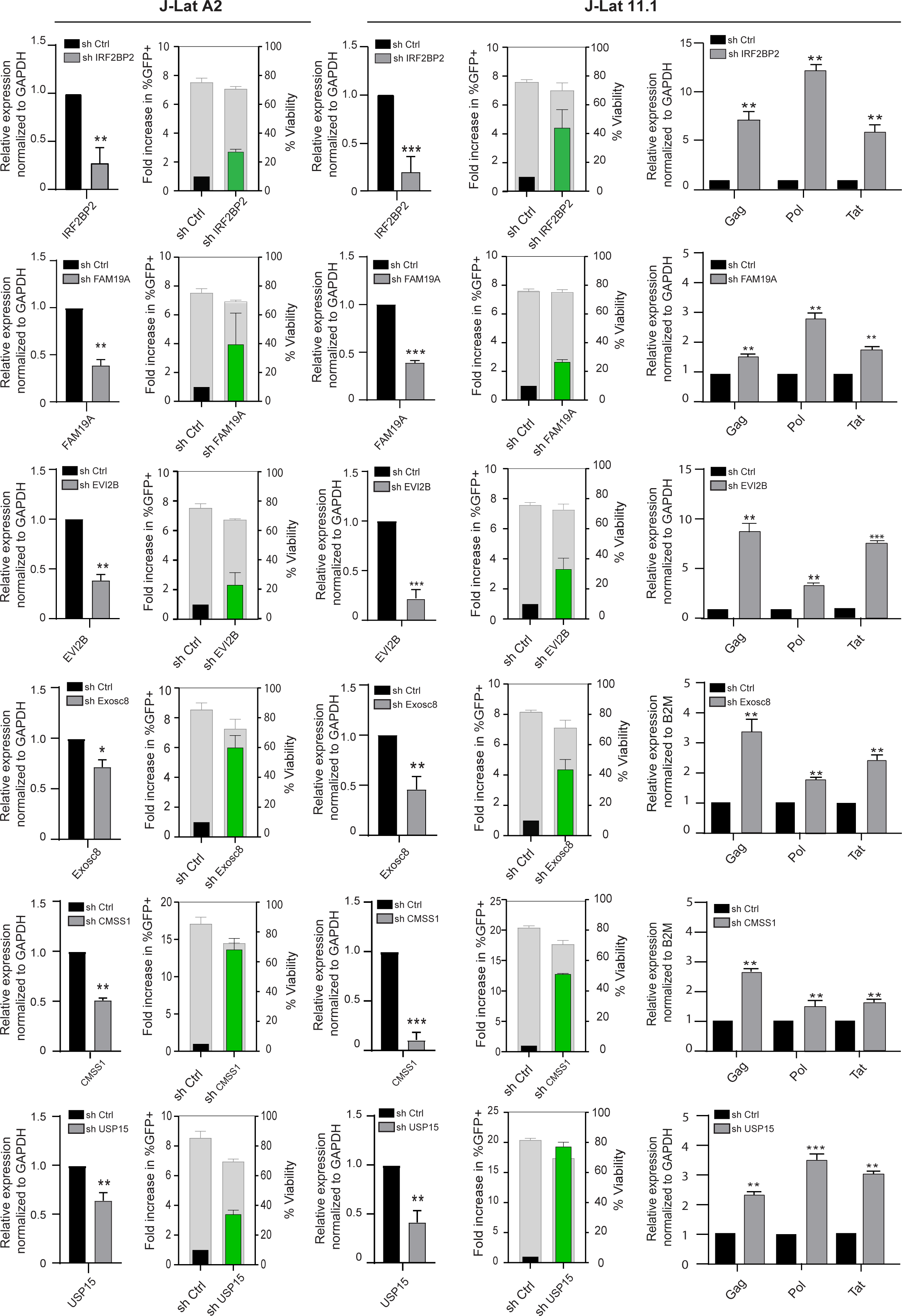
Functional validation of positive candidates. Re-activation of HIV-1 in J-Lat A2 and 11.1 cell lines was assessed by measuring the percentage of cells expressing GFP (green bars) and cell viability (gray bars) using flow-cytometry. Efficacy of knockdown by shRNA was quantitated in J-Lat 11.1 cells by RT-qPCR (left panel) as well as expression of viral genes Tat, Pol and Gag. RT-qPCR data (right panel) are presented as mean ± SD normalized to the control. Statistical significance was calculated using ratio-paired t-test and multiple comparison t-test on Log2 transformed fold changes (* – p < 0,05, ** – p < 0,01, *** – p < 0,001).

**Supplemental Figure 4.**
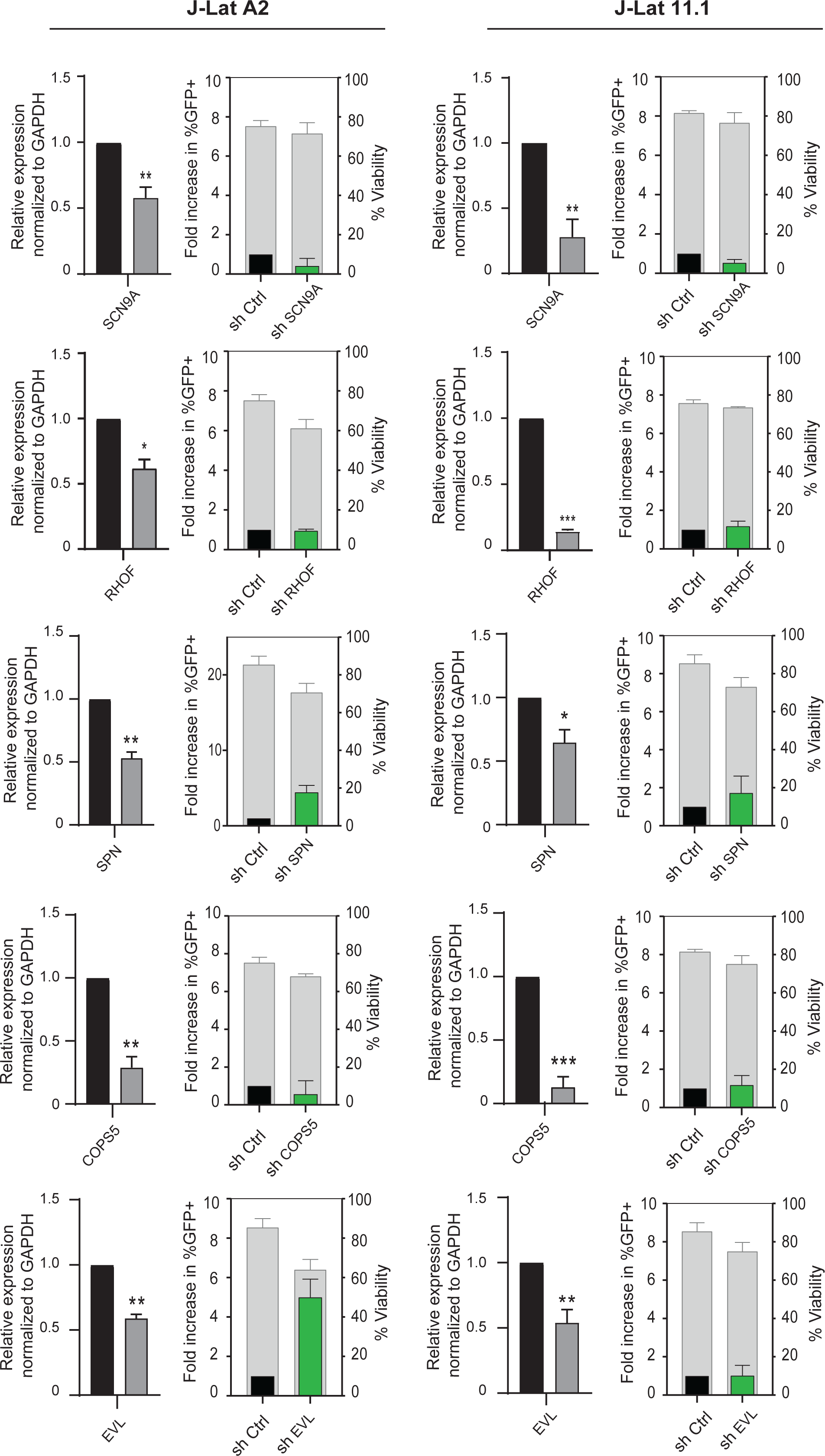
Functional validation of false positive candidates. Efficacy of knockdown by shRNA was quantitated by RT-qPCR. Re-activation of HIV-1 was assessed by measuring by the percentage of cells expressing GFP (green bars) and cell viability (gray bars). RT-qPCR data are presented as mean ± SD normalized to the control. Statistical significance was calculated using ratio-paired t-test and multiple comparison t-test on Log2 transformed fold changes (* – p < 0,05, ** – p < 0,01, *** – p < 0,001).

**Supplemental Figure 5.**
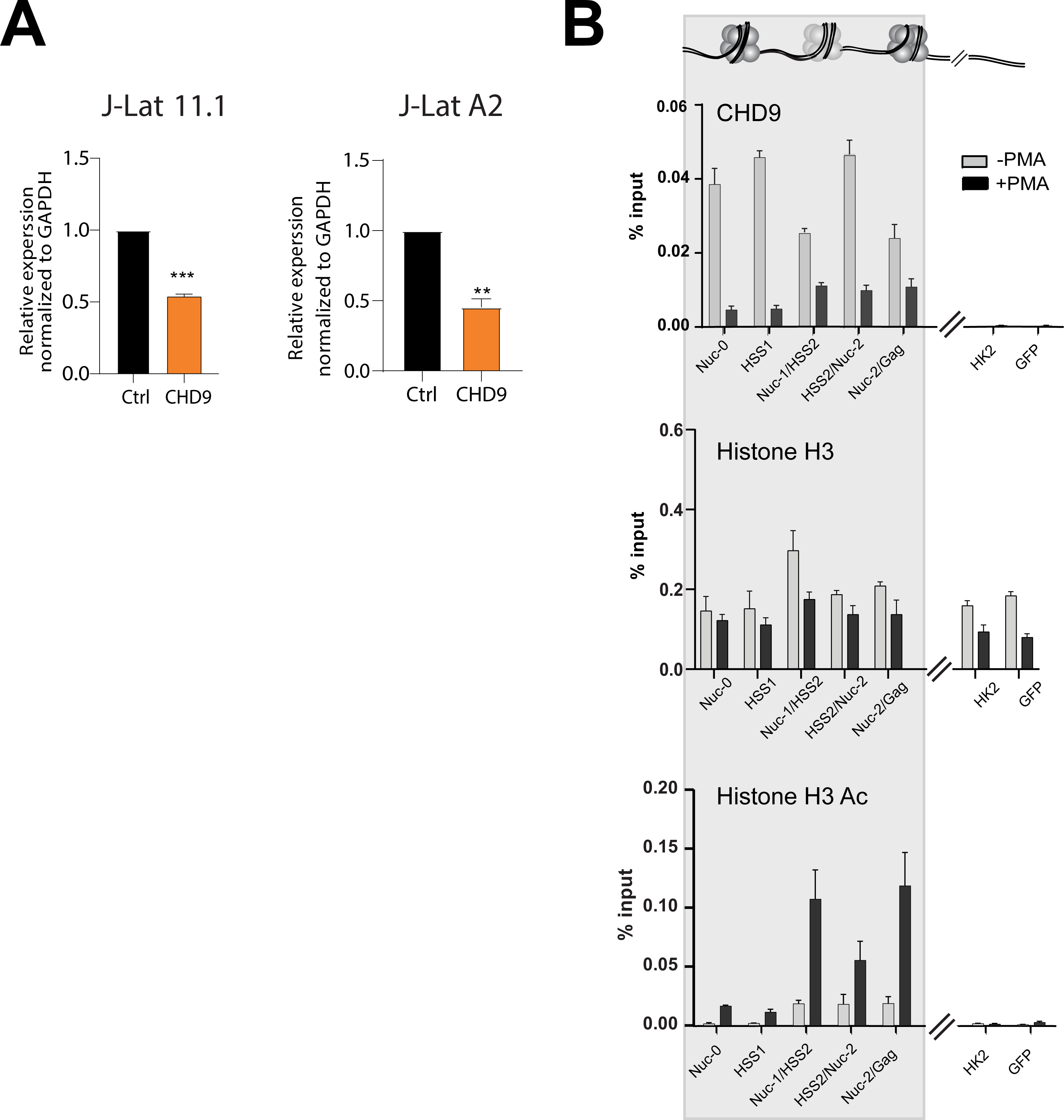
Additional data on the functional characterization of CHD9. (A) Efficacy of CHD9 knockdown in J-Lat A2 cells and J-Lat 11.1 cells by shRNA was quantitated by RT qPCR (B) Biological replicate of a ChIP-qPCR analysis of CHD9 binding to the HIV-1 5’LTR in untreated J-Lat 11.1 cells and PMA stimulated cells (Similar to data from figure 3G-H). Data are represented as percentage of the input and represent the average (±SD) of two technical replicates.

**Supplemental Figure 6.**
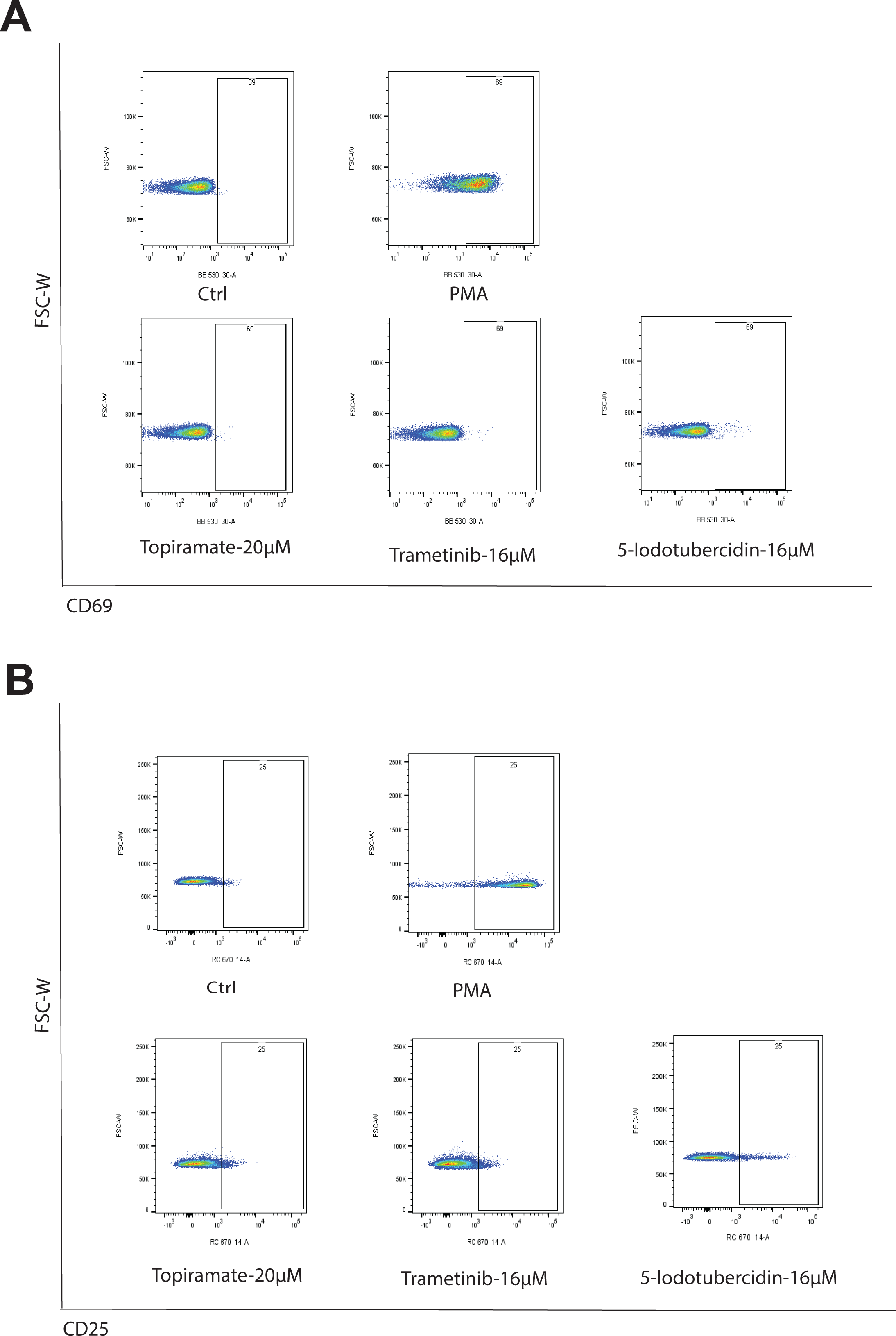
Characterization of T-cell activation by candidate LRAs. (A) FACS plot of staining for T-cell activation marker CD69 after treatment of primary CD4+ T cells from 6 healthy donors with candidate LRAs (5-Iodotubercidin, Topiramate and Trametinib). Treatment with PMA/Ionomycin PMA is used as a positive control (Plots correspond to data from figure 4F). (B) FACS plot of staining for T-cell activation marker CD25 after treatment with candidate LRAs (Plots correspond to data from figure 4G).

**Supplemental Figure 7.**
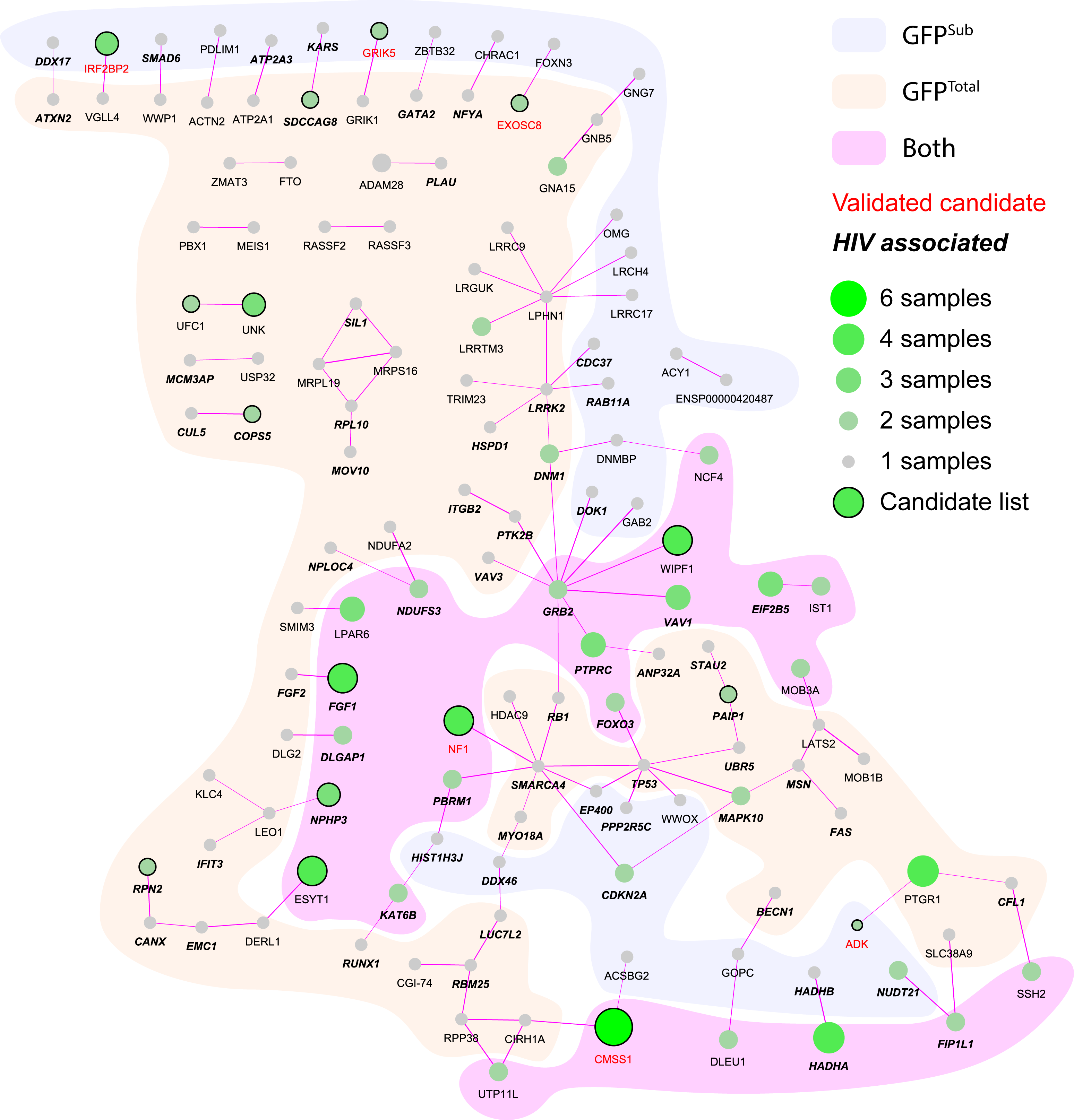
STRING analysis of protein coding candidate genes. STRING analysis indicates that many of the 598 protein coding candidate genes with an LOF score of 3 and higher from the GFP^Total^ and the GFP^Sub^ populations functionally interact. Validated candidate genes are indicated in red. Circle size and green shade indicate the number of samples the Gene Trap target gene is found in. Circled genes are part of our candidate genes list. Background color indicates if the Gene Trap target gene is found in the GFP^Total^, in the GFP^Sub^ or in both populations.

